# Metastatic tumor cells in bone marrow differ from paired neuroblastoma tumor and contain subsets with therapy-resistant characteristics

**DOI:** 10.1101/2024.09.13.612231

**Authors:** Caroline Hochheuser, Arjan Boltjes, Kaylee M. Keller, Simon Tol, Marieke van de Mheen, Carolina Pita Barros, Zeinab van Gestel-Fadaie, André B. P. van Kuilenburg, Sander van Hooff, Carlijn Voermans, Jan J. Molenaar, Godelieve A. M. Tytgat, Ilse Timmerman

**Affiliations:** Princess Máxima Center for Pediatric Oncology, Utrecht, the Netherlands; Department of Hematopoiesis, Sanquin Research and Landsteiner Laboratory, Amsterdam University Medical Centers, University of Amsterdam, Amsterdam, the Netherlands; Department of Research Facilities, Sanquin Research and Landsteiner Laboratory, Amsterdam University Medical Centers, University of Amsterdam, Amsterdam, the Netherlands; Laboratory Genetic Metabolic Diseases, Cancer Center Amsterdam, Amsterdam University Medical Centers, Vrije Universiteit Amsterdam, Meibergdreef 9, 1105 AZ, Amsterdam, The Netherlands; Department of Hematology, Amsterdam University Medical Centers, University of Amsterdam, Amsterdam, the Netherlands; Department of Pharmaceutical Sciences, University Medical Center Utrecht, University of Utrecht, Utrecht, the Netherlands; Department of Genetics, University Medical Center Utrecht, University of Utrecht, Utrecht, the Netherlands

**Keywords:** Bone marrow, metastases, intra-tumor heterogeneity, single-cell RNA sequencing, dormancy, therapy resistance, neuroblastoma, GD2

## Abstract

Bone marrow (BM) is a common site for solid tumor metastasis, often causing poor outcome. Here, we define the characteristics of BM-disseminated tumor cells (DTCs) using neuroblastoma as a model. We combined single-cell RNA-sequencing (scRNA-seq) and cell-surface protein analysis using 7 paired BM and primary tumor (PT) samples and found that DTCs contain a higher percentage of cycling cells and higher expression of neurodevelopmental genes compared to corresponding PT cells. In 6 patients, the copy number variation profile differed between PT cells and DTCs, indicating spatial heterogeneity. Within the BM, we detected dormant DTCs with potentially reduced chemosensitivity; this population contained cells expressing low levels of the immunotherapeutic antigen GD2 and increased NGFR expression. In conclusion, we characterized DTCs that are particularly challenging to target, offering new avenues for developing therapeutic strategies designed to target all subpopulations within the highly complex metastatic site, thereby preventing the development of drug-resistant clones.

## Introduction

Solid tumors are typically difficult to cure once they have metastasized to the bone marrow (BM). A growing body of evidence suggests that tumor cells occupy a specialized niche within the BM, which facilitates tumor cell colonization and supports their long-term survival (*1*). The interaction between newly arriving tumor cells and this microenvironment is critical for the cells to adapt to their new surroundings and acquire the molecular features that decrease their susceptibility to therapy, for example by entering a prolonged dormant state (*1*, *2*). In this dormant state, the DTCs can evade treatment and the immune response, thereby leading to subsequent reactivation and relapse. Together with the inherent heterogeneity of tumors (*3*), these adaptations to the bone marrow metastatic site present major challenges with respect to treatment. Although the primary site of most solid tumors has been studied extensively at the molecular level, the features of BM metastases are poorly understood. To study the characteristics of these so-called BM-disseminated tumor cells (DTCs), we used neuroblastoma (NB)—a solid tumor that often metastasizes to the BM—as a model. NB is a pediatric tumor arising from progenitor cells of the sympathetic nervous system (*4*). Despite significant advances in treatment, more than 40% of children with high-risk NB experience relapse. This results in a long-term cure rate of less than 50% for patients with high-risk NB (*5*). Moreover, relapse often originates from DTCs in the BM (*6*); thus, effectively targeting these DTCs is essential for improving long-term outcome.

Despite evidence indicating a heterogeneous nature of NB tumors, all high-risk patients receive a standard first-line treatment consisting of induction chemotherapy, surgical resection, high-dose chemotherapy, autologous stem cell transplantation, and radiation (*7*). Post-consolidation therapy typically consists of antibody-based immunotherapy directed against the disialoganglioside GD2 (*8*). Inter-patient heterogeneity is an intrinsic property of NB tumors and is driven by gene mutations and chromosomal aberrations, such as *MYCN* amplification (*9*). Intra-patient heterogeneity includes tumor cell plasticity—defined as the interconversion between two cell states. *In vitro*, both adrenergic (ADRN) and mesenchymal (MES) NB cell states have been described, with the latter being chemoresistant and less differentiated (*10–12*). The existence of NB cells with MES features *in vivo* has been suggested based on studies of patient samples by quantitative polymerase chain reaction (qPCR) (*13*, *14*), bulk RNA sequencing (*15*), single-cell RNA sequencing (scRNA-seq) (*16*), and epigenetics (*17*). Moreover, spatial heterogeneity between metastases and the primary tumor contributes to intra-patient heterogeneity. In NB, spatial heterogeneity has been demonstrated at the DNA level (*18*, *19*) as well as by performing (pseudo-) bulk gene expression analysis of non-paired samples (*20*, *21*) and can include increased mitochondrial DNA–derived transcripts in DTCs. Recent scRNA-seq studies of BM metastases of NB have shed light on tumor cell plasticity, interactions with immune cells (*20*, *22*), and evolutionary development of metastases in NB (*23*). However, the aspect of spatial heterogeneity remains poorly understood.

In this study, we generated a unique scRNA-seq dataset comprised of seven paired sets of BM-derived DTCs and PT cells in order to examine how metastatic tumor cells differ from the cells in the PT from which they originated, thereby eliminating the confounding factor of inter-patient heterogeneity. To effectively lower the risk of relapse and improve patient outcome, all DTC subpopulations in BM must be targeted. We therefore characterized the tumor cell subpopulations in the BM based on transcriptomics features and surface markers in order to examine the heterogeneity within the metastatic cell population and identify the DTC types that are most difficult to target. Our results reveal potential avenues for developing more effective treatment strategies targeted to metastatic disease and underscore the importance of considering intra-patient heterogeneity.

## Results

### Identification of tumor cells from primary tumor and bone marrow samples

First, we conducted single-cell transcriptomics analyses on PT samples and matching BM aspirates obtained from 7 patients with NB (Patients 1-7); BM aspirates (but no matching PT samples) were also obtained from 2 additional patients with NB (Patients 8 and 9). All samples were treatment-naïve diagnostic samples, and 4 of the 9 patients (Patients 2, 3, 8, and 9) had *MYCN* amplification (**Fig. 1A** and **Fig. S1A)**. Moreover, 8 patients had a subsequent event during their disease course. BM infiltration ranged from 0.4% to 80%; therefore, enrichment for DTCs was required by sorting CD34^neg^CD45^neg^CD90^pos^ and/or CD34^neg^CD45^neg^GD2^pos^ cells from the BM samples (see Methods). Simultaneously, the expression levels of additional NB and mesenchymal surface markers were recorded for each single DTC sorted into an individual well of the plate (“index sorting”), allowing us to link the cell’s transcriptome to the expression of 12 surface markers (**Fig. 1B** and **Fig. S1B-D**). PT and BM samples were single-cell sequenced using the SORT-seq protocol (*24*).

**Fig. 1.**
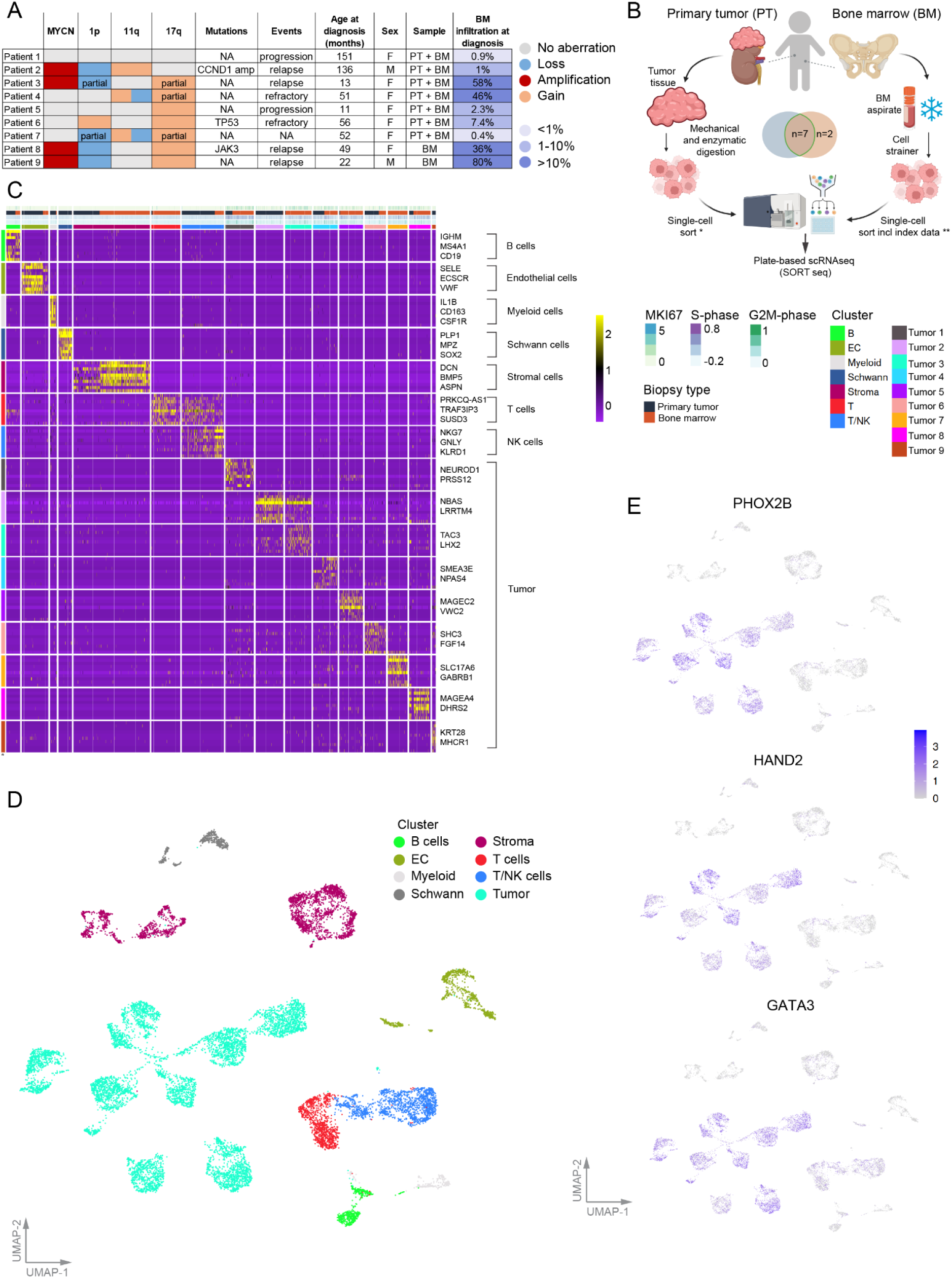
Cohort overview, workflow, and tumor cell identification. **A)** Clinical characteristics of the patient cohort at diagnosis. “Events” refers to the disease course for each patient. **B)** Schematic representation of the steps used to process fresh primary tumor samples and cryopreserved bone marrow aspirates. *All cells in the sample were sorted; **enrichment for CD34-CD45-CD90+ and/or CD34-CD45-GD2+ tumor cells (see representative gating strategy in Fig. S1C). **C)** Heatmap displaying the top 10 differentially expressed genes per cluster (see Fig. S1F). The cell type–specific markers used for annotation are indicated on the right. **D)** UMAP (uniform manifold approximation and projection) of the complete dataset with cell type annotation per cluster. **E)** UMAPs showing expression of the indicated NB marker genes in the adrenergic core regulatory circuit for all cells (tumor and non-malignant) shown in (D). BM, bone marrow; PT, primary tumor; NA, not available; EC, endothelial cells; NK = natural killer; CCND1 amp, cyclinD1 amplification; TP53, tumor protein 53; JAK3, Janus kinase 3.

We performed routine quality control checks on the sequencing data, including filtering on transcript number per cell and mitochondrial content (**Fig. S1E**), confounder removal, and correcting for differences in sample processing (for details, see Methods). Dimension reduction and clustering were then performed on the remaining 14,570 cells. Cell types were identified based on a comparison between gene expression patterns per cluster using the Human Cell Atlas (SingleR, **Fig. S1F-G**) and cell type–specific markers within the top 10 differentially expressed genes (DEGs) per cluster (**Fig. 1C**). Clusters with neuronal identity were initially identified as either tumor cells or Schwann cells (based on expression of SOX10 and S100B; data not shown) (**Fig. 1D** and **Fig. S1F-G**). Tumor cell identity was confirmed based on copy number variation (CNV) analysis using the inferCNV algorithm (**Fig. S1H**) and expression of the NB-specific markers *PHOX2B*, *HAND2*, and *GATA3* (**Fig. 1E**), which are part of the ADRN core regulatory circuit (*12*, *25*). A comparison of the bulk vs. single cell–sorted populations confirmed successful enrichment of tumor cells in the BM samples (**Fig. S1I**).

### Metastatic NB cells in the BM have higher cell cycle activity than their corresponding PT cells

Intra-patient heterogeneity—both spatial and temporal—can lead to diversification of the tumor’s clonal composition, thereby contributing to key events in disease progression (*18*, *19*, *26*). Given our unique access to 7 sets of paired diagnostic BM and PT samples, we aimed to identify this intra-patient heterogeneity between the primary and metastatic tumor cells at diagnosis. After selecting tumor cells from Patients 1-7, dimension reduction and clustering revealed patient-specific tumor populations (**Fig. 2A**, left panel). However, for each patient, the DTCs and PT cells formed distinct clusters (**Fig. 2A**, right panel), indicating both inter-patient and intra-patient heterogeneity. To characterize the differences we then performed differential gene expression analysis between all PT and DTCs from 7 patients combined, followed by gene set enrichment analysis (GSEA) (**Fig. 2B-D** and **Table S1**). Processes enriched in the PT cells included protein folding, autophagy, cell adhesion, epithelial-to-mesenchymal transition (EMT), apoptosis downregulation, and MAPK3 and VEGFA signaling (**Fig. 2C** and **Table S1**). In contrast, the most enriched processes in the DTCs were related to cell cycle status (**Fig. 2D** and **Table S1**). Consistent with this finding, the gene encoding the proliferation marker *MKI67* and genes involved in cell cycle regulation such as *TOP2A* and *MYBL2* (encoding DNA topoisomerase II alpha and MYB proto-oncogene like 2, respectively) were expressed at higher levels in the DTCs (**Fig. 2B, Table S1,** and **Fig. S2A**). To confirm the increased cell cycle status of DTCs, we categorized their cycling status as either high or low based on their G2M-phase and S-phase scores (**Fig. S2B**) and found that the BM cells in all 7 patients contained a higher percentage of cycling tumor cells compared to their corresponding PT cells (**Fig. 2E**).

**Fig. 2.**
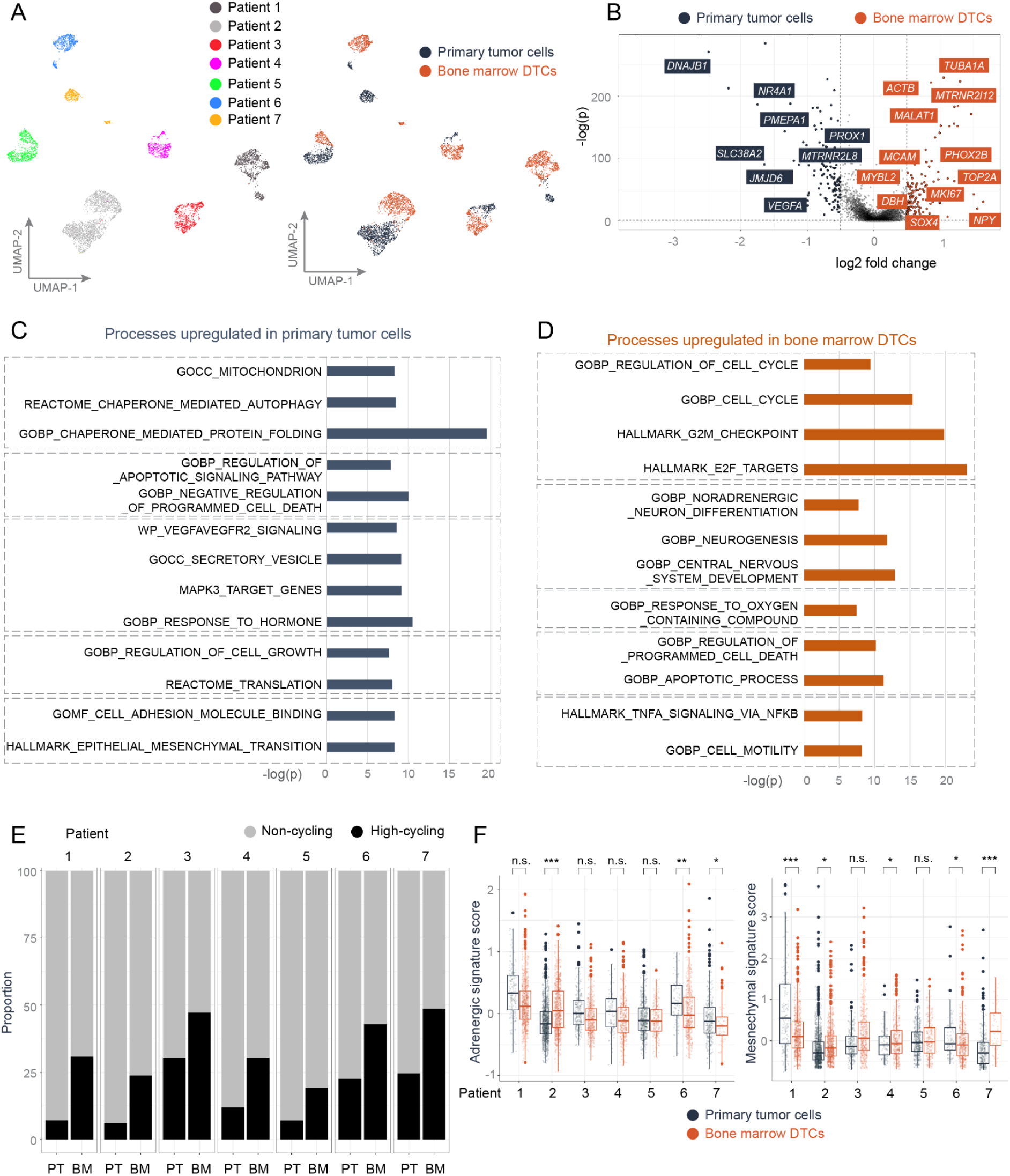
Differences in transcriptomics between primary tumor cells and paired metastatic cells in the bone marrow. Data from all 7 patients with paired primary tumor (PT) and bone marrow (BM) samples are included in these analyses. **A)** UMAP of tumor cells in patients 1-7 annotated by patient (left) or sample type (right). **B)** Volcano plot of differentially expressed genes between PT cells and BM-disseminated tumor cells (DTCs). Differential gene expression (DGE) was performed on all 7 PT samples vs. the corresponding 7 BM samples (see Table S1). **C-D)** Selected processes from the top-100 enriched gene sets in PT (**C**) and BM-DTCs (**D**) after gene set enrichment analysis (see Table S1 and Methods for details). **E)** Summary of the percentage of non-cycling and high-cycling tumor cells in PT and BM samples for each patient, based on the expression of G2M-phase and S-phase gene sets (see Fig S2B). **F)** Box plots summarizing the expression levels of ADRN (adrenergic) and MES (mesenchymal) signatures (*Gartlgruber et al, 2020*) (*17*) in PT and BM tumor cells for each patient. The boxes indicate the inter-quartile range, and the thick horizontal lines represent the median value. \**p*<0.05, \*\**p*<0.01, \*\*\**p*<0.001, and n.s., not significant (Wilcoxon rank sum test). Gene expression was assessed on scaled data in order to correct for processing differences between PT and BM samples (see Methods).

### Mesenchymal tumor cells are not enriched in bone marrow compared to the primary tumor

Next, we investigated the transition of NB cells between the ADRN state and the MES state, a process previously suggested to characterize a metastatic phenotype in NB cells (*27*) and which has been linked to therapy resistance and relapse (*11*, *12*, *17*). To determine whether at diagnosis the metastatic niche in the BM contains MES tumor cells, and to determine whether these tumor cells are enriched in the BM compared to the PT, we assessed the expression of published ADRN and MES signatures derived from both primary and relapsed NB tumors and from cell lines (*17*). The expression patterns of these signatures in paired PT and BM-DTCs varied among the patients of this cohort. Specifically, the MES signature expression was higher in the PT cells in one patient (Patient 1), but higher in the DTCs in another patient (Patient 7); in contrast, we found either no difference or only a small difference in the other 5 patients (**Fig. 2F**). Notably, we also found that ADRN and MES cells identified using other commonly used signatures derived solely from cell lines (*12*) yielded different results for some patients (**Fig. S2C**). In both cases, however, we found no overall enrichment of MES features in the metastatic cells at diagnosis. In fact, genes expressing markers for ADRN NB cells such as *PHOX2B* (paired like homeobox 2B), *DBH* (dopamine beta-hydroxylase), and *GAP43* (growth-associated protein 43) were among the higher expressed genes in DTCs, and genes involved in neuronal development such as *NPY* (neuropeptide Y), *STMN1* (stathmin 1), and *SOX4* (SRY-box transcription factor 4) were also highly expressed in DTCs (**Fig. 2B**, **Table S1**, and **Fig. S2A**). Consistent with these results, we found that gene sets such as “noradrenergic neuron differentiation” and “neurogenesis” were enriched in the metastatic DTCs (**Fig. 2D** and **Table S1**). Taken together, these findings indicate that at diagnosis DTCs are not predominantly MES cells, but rather display transcriptomics features characteristic for ADRN cells.

### Transcriptomics reveals heterogeneity in CNV profiles at diagnosis between DTCs and primary tumor cells

Genetic variation is a major contributor to tumor heterogeneity. To assess whether—and how—CNV profiles differ between paired sets of PT cells and DTCs, as well as the consequences of this difference for the transcriptome on a single-cell level, we measured gene expression intensity across chromosomes using inferCNV (*28*) (**Fig. S3**). We first used matching whole-genome sequencing/whole-exome sequencing (WGS/WES) data from the PT samples of 4 patients to confirm that inferCNV can detect the same and additional CNVs as DNA sequencing; a representative example of the WGS data from Patient 2 is shown in **Fig. S3C**. We found clear differences in chromosomal gains and losses between DTCs and PT cells for 6 of the 7 patients (**Fig. 3A** and **Fig. S3**, black arrows). For example, in Patient 1 chromosomal loss in chromosomes 10 and 11q was present in only a small percentage of PT cells, compared to nearly all DTCs in the same patient (**Fig. S3A**). Importantly, the 11q loss (which is commonly reported during routine diagnostics) was not indicated in the clinical information for Patient 1 (**Fig. 1A**), possibly because this aberration was present in only in a small percentage of PT cells. Conversely, for Patient 2 the majority of PT cells had a chromosome 9 loss, while the majority of DTCs in this patient lacked this chromosomal aberration (**Fig. 3B-C** and **Fig. S3B**). To gain insight into the biological consequences of these clonal differences, we performed differential gene expression analysis and GSEA between the aberrant and non-aberrant tumor cells, using the chromosome 9 CNV in the PT of Patient 2 as an example (**Fig. 3C-D** and **Table S2**). We found that processes related to migration and invasion, as well as TGFβ and TNFα/NF-κB signaling—both of which play a role in migration and EMT induction (*29*, *30*)—were downregulated in the cells with chromosome 9 loss (**Fig. 3D**). These findings provide evidence of spatial CNV heterogeneity in NB cells at diagnosis at the transcriptomic level.

**Fig. 3.**
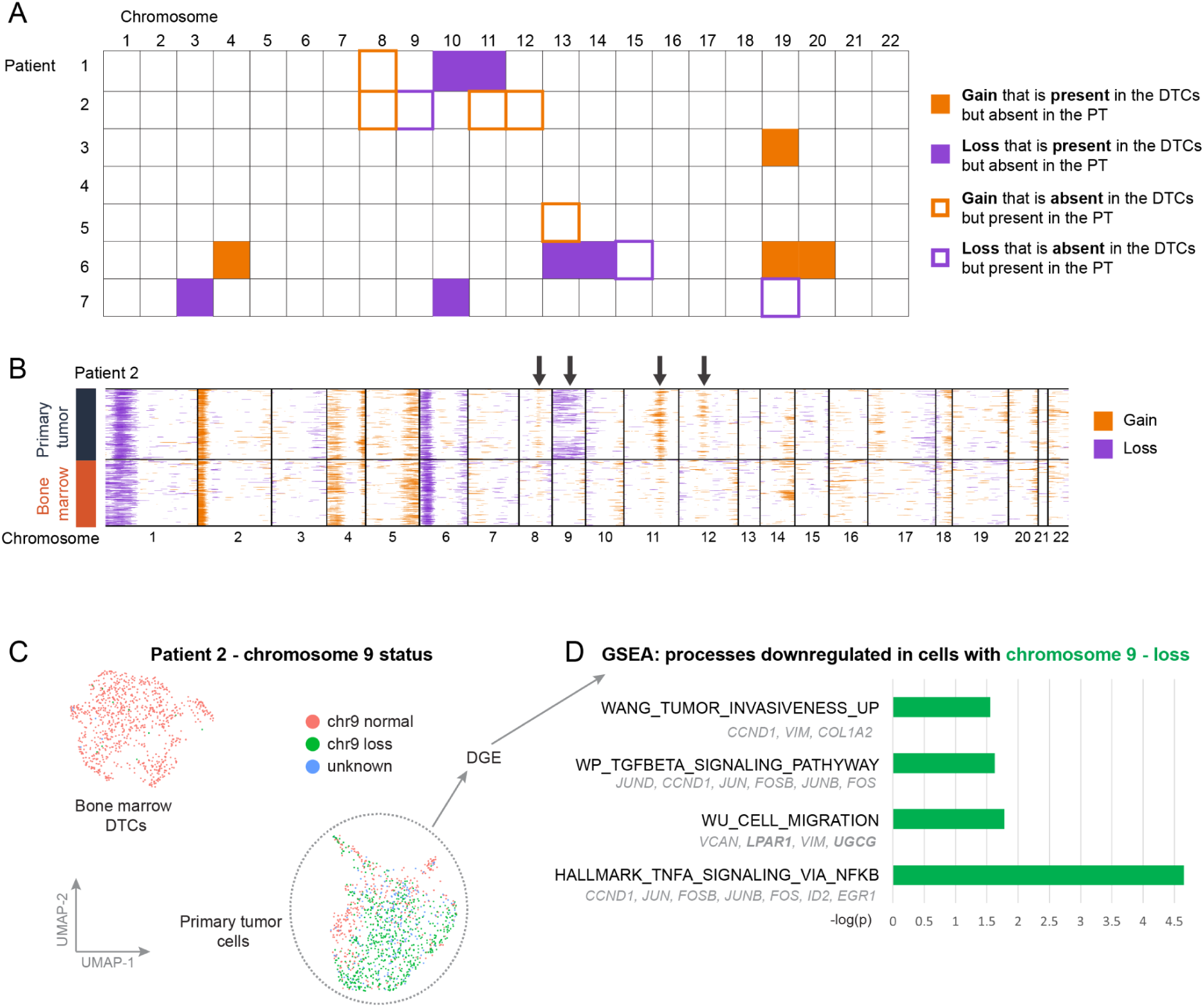
Heterogeneity in copy number variation between primary tumor cells and DTCs. **A)** Overview of heterogeneity in copy number variation (CNV) in the somatic chromosomes between PT and BM samples in Patients 1-7. Purple and orange indicate a chromosomal loss and gain, respectively. Full squares indicate aberrations that are present in the DTCs but absent in the PT, while open squares indicate aberrations that are present in the PT but are absent in the DTCs (see Fig. S3 for details). **B)** CNV plot (inferCNV) for Patient 2, depicting expression intensity across each somatic chromosome in PT cells and DTCs. The arrows indicate CNV heterogeneity between PT cells and DTCs. Purple and orange indicate a loss and gain, respectively. **C)** UMAP of tumor cells for chromosome 9 status in Patient 2; red, green, and blue indicate no loss/gain (“normal”), loss, and unknown respectively. DGE, differential gene expression. **D)** Selected processes involving the gene sets downregulated in cells with a loss in chromosome 9 in PT cells in Patient 2. A selection of differentially expressed genes is shown under each gene set (see also Table S2).

### Both cycling and dormant tumor cells with differing drug sensitivity coexist in the metastatic bone marrow population

Because recurrent disease often originates in the BM, identifying the detailed composition of the metastatic tumor cell population is essential in order to effectively target all subpopulations of tumor cells. Therefore, we comprehensively examined the transcriptomic and phenotypic heterogeneity of the metastatic DTC population at diagnosis. After integrating the DTCs of all 9 patients, we identified a subset of cells (clusters 1 and 4 in **Fig. 4A-B**) with high cell cycle activity (**Fig. 4A-B**). To determine whether this subset has different drug sensitivity compared to the other clusters, we examined the gene expression signatures associated with sensitivity to chemotherapeutic drugs (DepMap database) in the clusters. We found that the cycling clusters (i.e., clusters 1 and 4) scored higher in their predicted sensitivity to drugs commonly used in NB induction therapy such as vindesine and vincristine (**Fig. 4C** and **Table S3**). Next, we examined the transcriptomic characteristics of the cells within the other subsets (i.e., the cells with low cell cycle activity). The BM provides a protective niche for hematopoietic stem cells, maintaining a dormant state (*1*). To determine whether metastatic NB cells “hijack” this niche and become dormant—as hypothesized previously for a subset of metastatic tumor cells in breast cancer (*2*)—we measured the expression of published signatures that indicate tumor dormancy (*31*, *32*) in DTCs. We found that a small number of tumor cells within the non-cycling population scored positive for the combined dormancy signature (**Fig. 4D-E** and **Table S4**). Notably, these dormant cells were not detected in all patients’ DTC populations (**Fig. S4A**). Moreover, the signature scores for sensitivity to chemotherapeutics such as vindesine and vincristine were reduced in dormant cells, but also in other non-cycling cells (**Fig. 4F**); however, the signature scores for sensitivity to drugs such as etoposide and topotecan were reduced in the dormant population, compared to other non-cycling cells or proliferating cells (**Fig. 4F**). The dormant cells—and the other non-cycling cells—were characterized by their increased expression of genes related to the TNFα/NF-κB signaling pathway (**Fig. S4B**), a pathway known to help protect against apoptosis (*33*). Notably, expression of the dormancy signature was not exclusive to DTCs, but was also present in PT cells (**Fig. S4C-D**). Thus, within the DTC population in treatment-naïve BM samples, cells with increased cell cycle activity coexist with cells that exhibit dormancy features and a signature consistent with reduced drug sensitivity.

**Fig. 4.**
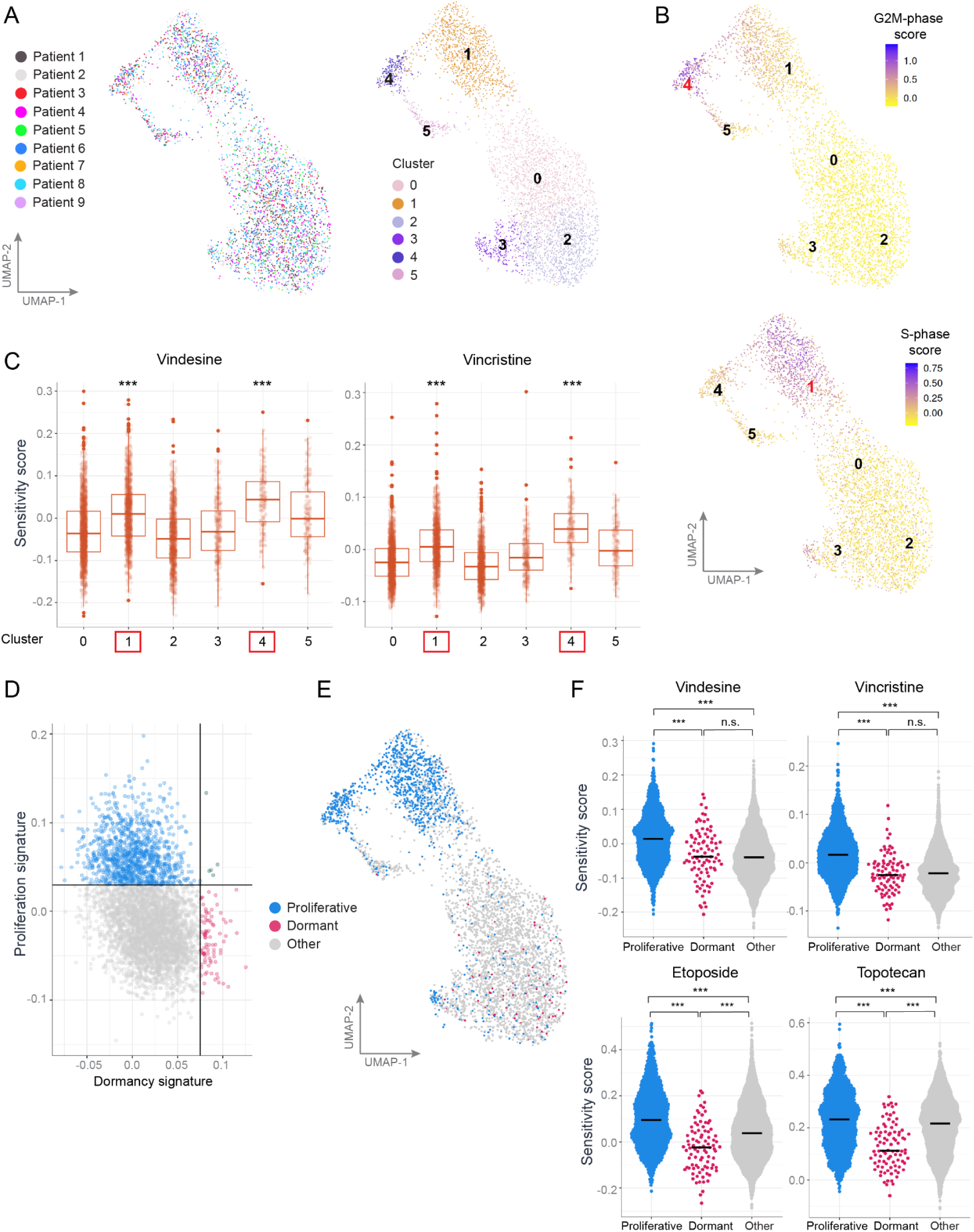
Drug sensitivity, cell cycle status, and dormancy features in DTCs in diagnostic bone marrow samples. **A)** UMAP and clustering of BM-DTCs from Patients 1-9 after integration. **B)** G2M-phase (top) and S-phase (bottom) scores indicate cycling cells within the BM DTC population, primarily in cluster 4 and cluster 1 in G2M-phase and S-phase, respectively. **C)** Box plots summarizing expression of gene signatures associated with sensitivity to vindesine (left) and vincristine (right) from DepMap database, for clusters 1-5 (see Table S3 for other drugs used in NB induction therapy). Boxes indicate the inter-quartile range, and the thick horizontal lines indicate the median value. Clusters 1 and 4, the high-cycling clusters based on G2M-phase and S-phase scores, respectively, are highlighted in red boxes. \*\*\**p*<0.001 versus the remaining clusters, except for the comparison between cluster 1 and cluster 5, which was non-significant. Kruskal-Wallis rank sum test followed by Dunn’s test for multiple comparisons). **D)** Expression of a dormancy gene signature (constructed from the overlap between 2 published signatures (*31*, *32*); see Table S4) was used to categorize DTCs as dormant, proliferating or neither (‘other’). **E)** UMAP showing distribution of dormant and proliferating DTCs in all 9 patients. **F)** Violin plots summarizing the drug sensitivity signatures to vindesine, vincristine, etoposide, and topotecan (derived from the DepMap database) in dormant and proliferating cells (see Table S3 for the complete list of scores for all chemotherapeutic drugs in dormant and proliferating cells). The thick horizontal lines indicate the median value. \*\*\**p*<0.001 and n.s., not significant (Kruskal-Wallis rank sum test followed by Dunn’s test for multiple comparisons).

### Linking transcriptomics and surface marker expression reveals characteristics of tumor cells with low GD2 expression in the bone marrow

Next, we integrated our scRNA-seq data with our analysis of cell-surface marker expression—including GD2 (**Fig. S1B**)—obtained by performing flow cytometry during cell sorting. GD2 is an NB marker shown to be clinically important for diagnostic purposes and anti-GD2 immunotherapy (*34*, *35*). Heterogeneity of GD2 expression in tumor tissues has been described and can affect both the detection and targeting of primary and metastatic tumor cells (*36–38*). We therefore sought to identify the characteristics of GD2-low DTCs and examine alternative approaches to detect these cells. We found that the surface expression of GD2 was not correlated with the mRNA levels of the two genes most relevant to GD2 synthesis (**Fig. S5A),** emphasizing the importance of measuring GD2 surface levels. Flow cytometric analysis of the BM samples in Patients 1-9 (**Fig. 5A**) revealed that 93.7% of DTCs had either medium or high levels of GD2 expression (**Fig. 5B**); however, in 3 out of 9 patients GD2 expression varied widely, and GD2 expression was either absent or low in 6.3% of DTCs (**Fig. 5A-B**). Importantly, tumor cell identity was confirmed using linked scRNA-seq data (**Fig. 1**, **Fig. S1H**, and **Fig. S5B**). An analysis of surface protein levels of the alternative NB markers CD56 (also known as neural cell adhesion molecule, or NCAM), and CD90 (also known as the Thy-1 cell surface antigen) showed that within the GD2-neg/low DTC population, 86.4% and 85.3% of cells were positive for NCAM and Thy-1, respectively (**Fig. 5B**). Because we enriched for DTCs by gating on Thy-1 and/or GD2, cells lacking both of these markers would be missed in our analysis; however, based on a pilot study, the percentage of Thy-1-neg/GD2-neg DTCs was extremely small (**Fig. S1D**).

**Fig. 5.**
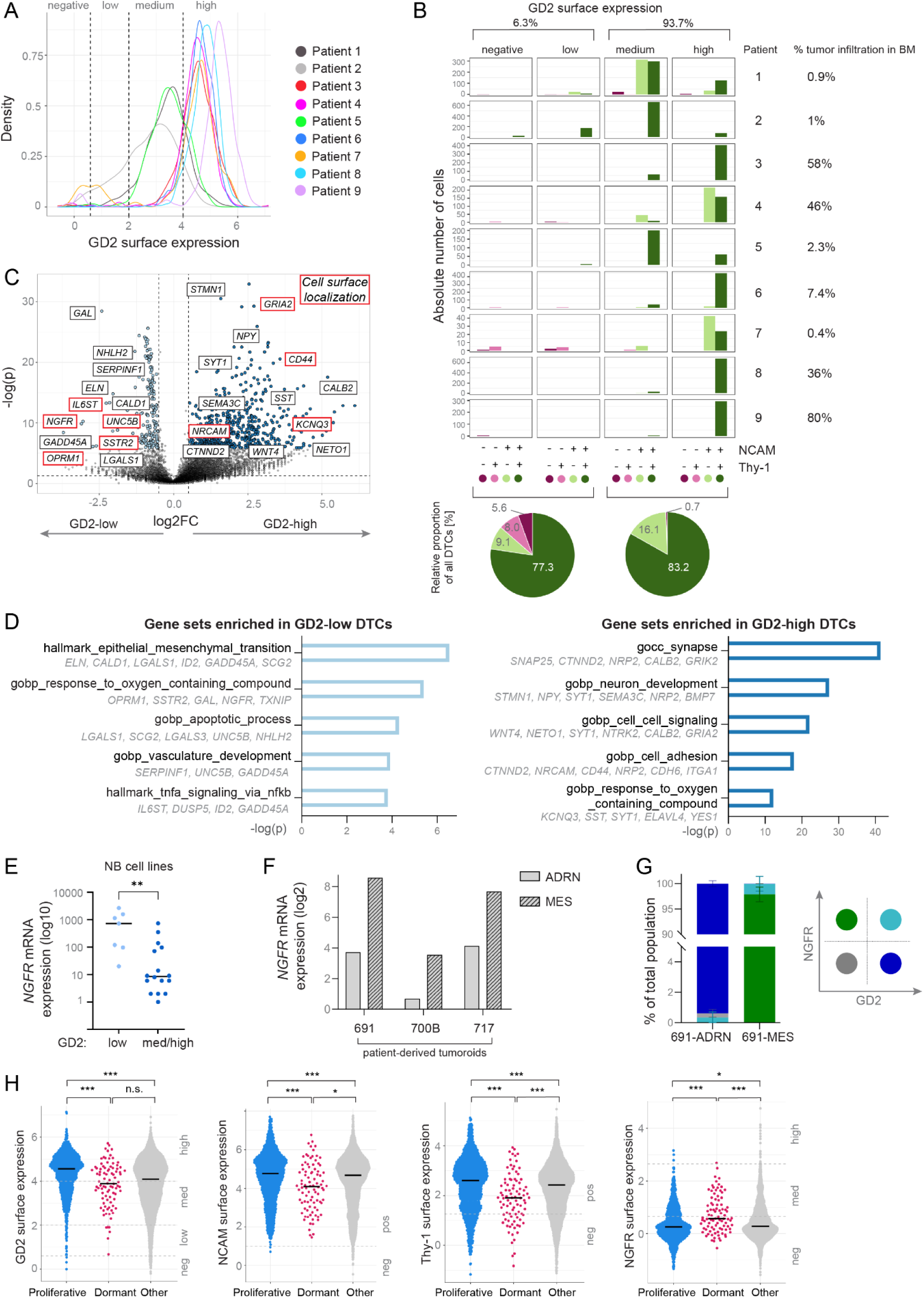
Linking surface protein expression to transcriptomics patterns in disseminated tumor cells. **A)** Density plot of relative surface expression of GD2 in DTCs from the BM of all 9 patients. The thresholds for expression intensity (negative, low, medium, or high GD2 expression) are indicated and are based on the distribution of each patient’s DTC population in relation to non-malignant cell clusters (see Fig. S5E). **B)** Bar graphs indicating absolute number of tumor cells in each GD2 category expressing the alternative NB surface markers NCAM (CD56) and/or Thy-1 (CD90). Shown below are pie charts depicting the percentage of each population in the respective GD2 expression categories (date are based on all 9 patients combined). **C)** Volcano plot showing differentially expressed genes (DEGs) between GD2-neg/low and GD2-high DTCs in Patients 1 and 2, the two patients who had the largest variation in GD2 expression. Genes indicated in red boxes are annotated in UniProt as encoding cell-surface proteins (see Table S5). **D)** Selected processes from the top-50 enriched gene sets in GD2-high and GD2-neg/low cells after GSEA. A selection of DEGs in each gene set is shown (see Table S5 and Methods for details). **E)** Summary of *NGFR* mRNA levels measured in 23 NB cell lines, categorized into GD2-low or GD2-medium/high groups based on GD2 levels determined by lipidomics. The thick horizontal lines indicate the median value; note that the *y*-axis is a log scale. Experiments were performed in triplicate. \*\**p*<0.01 (Mann-Whitney U test). **F)** Summary of *NGFR* mRNA (log2) measured in three pairs of ADRN (light gray bars) and MES (hatched bars) tumoroids derived from a public microarray gene expression dataset (GSE90803). **G)** Bar graph summarizing the percentage of GD2+ (dark blue), NGFR+ (green), double-positive (light blue), and double-negative (grey) 691-ADRN and 691-MES tumoroid cells. Surface expression was measured using flow cytometry, and error bars indicate the SD (n=3 independent experiments). **H)** Violin plots summarizing surface expression of GD2, NCAM, Thy-1, and NGFR measured in DTCs that are categorized as dormant, proliferating or neither (‘other’) (defined as shown in Fig. 4d). \**p*<0.05, \*\**p*<0.01, \*\*\**p*<0.001, and n.s., not significant (Kruskal-Wallis rank sum test followed by Dunn’s test for multiple comparisons).

To compare between GD2-neg/low and GD2-high DTCs, we performed differential gene expression analysis and subsequent GSEA, focusing on DTCs in Patients 1 and 2, as these showed the largest variation in GD2 expression (**Fig. 5C-D** and **Table S5**). Gene sets enriched in the GD2-neg/low DTCs were related to TNFα signaling via NF-κB, and EMT. In contrast, the gene sets enriched in the GD2-high DTCs were related to cell adhesion and neuronal development. To screen for potential alternative surface markers specific to GD2-low DTCs, we identified the DEGs that encode surface proteins based on publicly available subcellular localization data (see Methods). We found that *NGFR*, which encodes the nerve growth factor receptor (also known as p75^NTR^ or CD271), was among the genes that were expressed at high levels in GD2-low DTCs (**Fig. 5C**). In a fraction of GD2-low DTCs we measured relatively high surface expression of NGFR, based on flow cytometric analysis of BM samples (**Fig. S5C**). To support these findings, we performed additional experiments using cell lines and patient-derived tumoroids. First, we performed lipidomics and transcriptomics analyses on a set of 23 NB cell lines and found that cell lines that express low levels of GD2 have higher levels of *NGFR* expression (**Fig. 5E)**, and this difference was particularly apparent in non-*MYCN*–amplified (MNN) cell lines (**Fig. S5D**). Furthermore, our analysis of patient-derived tumoroid pairs—performed using publicly available microarray gene expression data—revealed higher expression of *NGFR* in mesenchymal tumoroids compared to adrenergic tumoroids (**Fig. 5F**). We then confirmed that the increased *NGFR* mRNA levels translate to higher surface levels of NGFR in the mesenchymal cells of the 691-tumoroid pair and coincided with low surface levels of GD2 (**Fig. 5G**).

To assess whether NGFR and the NB markers GD2, NCAM, and/or Thy-1 can detect dormant tumor cells in BM, we compared surface expression of these proteins between dormant and proliferating DTCs. GD2 was lower expressed in non-proliferative compared to proliferative cells and the dormant DTC population contained GD2-low cells. Dormant DTCs had significantly lower surface levels of NCAM and Thy-1; conversely, we found higher surface levels of NGFR in dormant DTCs compared to both proliferating DTCs and other non-proliferative DTCs (**Fig. 5H**). Taken together, integration of scRNA-seq and cell-surface protein analyses unraveled characteristics of GD2-low DTCs, including increased expression of MES genes and increased surface expression of NGFR. Dormant subpopulations of DTCs, which exhibited reduced predicted sensitivity to chemotherapy, contained cells expressing GD2 at low levels and showed increased surface expression of NGFR.

## Discussion

Currently, assessing risk and decision-making regarding therapeutic interventions for high-risk NB are based largely on the characteristics of the primary tumor. However, the common site of metastatic disease in NB—the bone marrow—poses a large risk for disease progression and relapse and thus requires more specific and effective targeting. In this study, we therefore investigated tumor cells that metastasized to the BM, both in comparison to the PT as well as with respect to heterogeneity within the DTC population. Our key findings include differences in cycling status and CNV profiles between DTCs and the matched PT cells, as well as identification of DTC subpopulations with characteristics that likely render them less susceptible to current treatment strategies.

Comparing DTCs to PT cells from matched samples, we found a higher percentage of cycling tumor cells in the BM compared to the PT. This finding is consistent with a previously reported study involving 4 patients with NB, which showed higher expression of a cell cycle–related gene set in metastatic cells in the BM compared to circulating tumor cells (*39*). High cycling activity may be linked to more aggressive features of tumor cells, as observed in other cancer types (*40*). Additionally, mesenchymal characteristics are suggested to be associated with tumor aggressiveness, as well as therapy-resistance and metastasis in various solid tumors, including NB (*11*, *12*, *23*). However, applying two commonly used MES signatures (*12*, *17*) did not reveal a consistent trend of mesenchymal-like phenotype enrichment in metastatic DTCs compared to PT cells from diagnostic samples of patients with NB. We also found no enrichment of the ADRN signature in either PT samples or DTCs; however, differential gene expression and GSEA did show transcriptional features consistent with ADRN NB cells—namely, high expression of *PHOX2B* and *DBH*—in the DTCs. These findings are consistent with findings reported recently by Fetahu and colleagues (*20*), who described a similar differentiation status between metastatic cells and primary NB (albeit unpaired), and a primarily adrenergic phenotype in metastatic cells. Overall, these findings suggest a predominantly ADRN phenotype for metastatic NB cells in the BM niche. Previous studies hypothesized that an enrichment of MES characteristics is to be expected primarily during the process of metastasis in circulating tumor cells (*39*) or in response to chemotherapeutic pressure (*12*, *16*, *17*, *41*); thus, identification and characterization of MES NB cells in the peripheral blood and/or post-treatment BM samples may reveal new options for targeting this subpopulation.

Our data also provides compelling evidence of spatial CNV heterogeneity between metastatic and PT cells at diagnosis at a transcriptomics level, suggesting that clonal evolution events occur during the early steps of tumor development and/or during metastasis. These findings extend our current knowledge regarding intra-patient heterogeneity in NB, which stems from genomics studies comparing the PT between diagnosis and relapse(*19*) or between the PT and liquid biopsies (*26*), as well as relatively few studies that included metastatic samples (*18*, *42*). Together with our findings, these studies underscore the urgent need to gain new insights into how the CNV profile differs between metastatic cells and the PT. Combining BM aspirates with PT material for CNV analysis during routine diagnostics could therefore provide a valuable tool for accurately assessing risk and guiding the clinical decision-making process (*18*, *43*). Linking CNVs to changes in transcriptomics enabled us to investigate how these CNVs confer either more aggressive or less aggressive features to the tumor subpopulations. For example, the PT cells with chromosome 9 loss in Patient 2 were characterized by decreased migratory capacity, suggesting reduced potential for these cells to invade the BM. Loss of genes that are important for tumor cell migration may contribute to the less aggressive nature of DTCs with chromosome 9 loss. For example, the *LPAR1* gene located on chromosome 9q was previously shown to be required for the metastasis of melanoma cells (*44*). Such insights into the biological implications are valuable for understanding the consequences associated with CNV heterogeneity. More complex approaches that integrate scRNA-seq and scDNA-seq (reviewed and further developed by Edrisi *et al*. (*45*)) can provide an even more in-depth analysis of clonal evolution in relation to the clones’ transcriptome.

Our study further reveals heterogeneity among DTCs in the BM metastatic niche. For example, we detected cells within the non-cycling DTC clusters that exhibited a dormancy transcriptomics signature derived from previous studies involving lung, colorectal, and breast cancer (*31*, *32*) that primarily used a lack of proliferation for identifying dormant cells. In our data, the characteristics of dormancy were detected in only a subset of non-cycling DTCs, suggesting that these DTCs are characterized by more than simply a lack of proliferation. The next step toward improving the definition of this critical cell population—and the features that distinguish these cells from other non-cycling cells—is to establish these signatures in NB models and incorporate other features of tumor cell dormancy, for example metabolic changes, increased autophagy, or an increase in the p38/ERK ratio. Importantly, we found that dormancy coincided with a reduction in the cells’ predicted drug sensitivity—particularly to etoposide and topotecan—suggesting reduced chemotherapy-mediated eradication of this subpopulation. The dormant cells—as well as other non-proliferating cells—also had high expression of genes involved in TNFα signaling via NF-κB, a trait recently attributed to so-called “persister cells” (i.e., cells that survive chemotherapy) (*46*). The fact that dormant cells were detected—albeit in small numbers—in treatment-naïve diagnostic samples suggests that mechanisms other than chemotherapeutic pressure may induce cellular dormancy in NB DTCs (*47*). These mechanisms may include acidification via hypoxia (*48*), as well as signaling pathways inherent to perivascular (*49*) and endosteal hematopoietic stem cell (HSC) niches (*50*), including TGF-β signaling (*51*).

By linking flow cytometry and sequencing data at the single-cell level, we studied heterogeneity in DTCs in detail. With respect to the immunotherapeutic target antigen GD2, we found that surface protein levels—rather than mRNA levels of GD2-synthesizing enzymes—were essential to our analysis, in contrast to previous studies (*52*, *53*). Variations in GD2 surface expression, which have also been reported in a few other studies (*36–38*), likely have important biological and clinical implications, as reduced GD2 expression was shown to diminish the efficacy of NK cell–mediated killing *in vitro* (*37*) and has been suggested as a mechanism by which tumor cells acquire resistance to immunotherapy (*54*, *55*). Not surprisingly, low GD2 expression has therefore been associated with relapse in patients receiving anti-GD2 immunotherapy (*37*). The presence of such GD2-low/neg cells in treatment-naïve samples suggests that NB cells with intrinsically low GD2 surface expression exist, or that factors other than immunotherapeutic pressure can decrease GD2 expression. In light of recent developments regarding anti-GD2 treatment during induction therapy (*56*) and alternative therapies targeting GD2 (*35*, *57*), investigating strategies designed to increase GD2 expression is particularly interesting (*58*) and should be complemented by a search for additional surface markers. To address this, we examined the surface marker and gene expression patterns in GD2-low/neg DTCs and found that approximately 85-86% of these cells express the NB markers Thy-1 and/or NCAM. A previous study by Lazic et al. found similar variations in the expression of NB surface markers using multiplex imaging (*38*). Thus, a multi-marker approach should be used to capture the heterogeneity of DTCs, and this approach should include markers that can discriminate between tumor cells and mesenchymal stromal cells (*59*).

Our characterization of the transcriptome in GD2-low versus GD2-high DTCs confirmed a more MES-like phenotype, as suggested previously (*55*) based on increased expression of *ELN*/*CALD1*/*LGALS1*, genes which have been linked—either directly or indirectly—to the induction of EMT (*60–62*). Importantly, the dormant DTC population also contained GD2-low cells, which would be difficult to target with the standard treatments currently used in clinical practice; specifically, in addition to being less susceptible to chemotherapy due to their reduced cell cycle activity, these cells may also evade anti-GD2 immunotherapy. Identifying alternative surface markers for these dormant GD2-low DTCs with MES characteristics is therefore essential for improving patient outcome. Here, we identified NGFR as a potential additional surface marker for detecting a subset of MES-like GD2-low DTCs, as well as a subpopulation of dormant DTCs. Based on previous findings in melanoma (*63*, *64*), NGFR may be functionally involved in inducing a low-proliferative state of DTCs and may therefore confer treatment resistance. Specifically, Restivo and colleagues found that the high NGFR levels in metastatic cells had to return to basal levels in order for the cells to regain their proliferative capacity (*63*). Interestingly, Lehraiki et al. showed that both NGFR upregulation and drug resistance are induced by NF-κB signaling stimulated by TNFα (*64*). Notably, our data identified this signaling axis as a promising avenue of study, as the TNFα/NF-κB gene set was enriched in DTCs versus PT cells at diagnosis, particularly in non-cycling and GD2-low DTC subpopulations. Findings from other studies suggest that TNFα/NF-κB signaling contributes to EMT, protection against apoptosis, and drug resistance (*29*, *33*, *46*), and therefore also plays a role during treatment. Targeting the GD2-low, MES, dormant DTC subpopulations as effectively and early as possible based on their quiescence, surface marker profile, and other biological features, will likely be the key to preventing the outgrowth of resistant clones and relapse.

In summary, our comparison between paired sets of primary and metastatic NB cells—without the possible confounding effects of inter-patient differences—reveals that DTCs differ from the PT cells both in their cycling activity and CNV profiles. Our in-depth analysis of DTCs supports the notion of a highly complex metastatic population that will likely require several complementary treatment strategies in order to efficiently eliminate all metastatic subpopulations. Taken together, these findings provide a clearer understanding of the metastatic niche in the BM and provide a starting point for further analyses regarding transcriptomic and phenotypic plasticity, and the persistence of subpopulations during treatment and in relapse. Studying these dynamics within a larger patient cohort covering all major genetic subgroups and including an extended panel of surface markers will likely facilitate the characterization and targeting of minimal residual disease and improve outcome, particularly in high-risk patients.

## Materials and Methods

### Study design and sample processing

This study was designed to investigate the transcriptomic differences between primary and metastatic tumor cells derived from the same patient. To this end, we collected matched samples from tumor biopsies and bone marrow aspirates. For further investigation of the BM DTC population, additional (non-matched) BM samples were included. All samples were obtained at diagnosis, prior to starting chemotherapy. After routine diagnostic procedures, the remaining material was used if written informed consent was provided by the patient and/or parents. The use of these samples was reviewed and approved by the biobank committee of the Princess Máxima Center (PMCLAB2019.067). Primary tumor and bone marrow samples of 7 patients and 9 patients, respectively, with high-risk NB were included in this study (the clinical characteristics are summarized in **Fig. 1A**). Bone marrow infiltration at diagnosis was determined by qPCR (*14*).

### Primary tumor samples

Primary tumor samples were processed fresh within 4 hours after surgery. The tissue was first cut into pieces <1 mm^3^ with a scalpel, and subsequently digested with collagenase type I, II and IV (2.5 mg/ml each, Life Technologies) for up to 1 hr at 37°C. The samples were then filtered through a 70-μm cell strainer, washed with DMEM (centrifuged at 300 g for 10 mins at 4°C) and further dissociated into single cells using the NeuroCult Chemical Dissociation Kit (STEMCELL Technologies; #05707) in accordance with the manufacturer’s instructions. The cells were then washed and stained with DAPI (Merck; #D9542) and DRAQQ5 (BioLegend; #424101) at 4°C, and live cells (DAPI-negative/DRAQ5-positive) were sorted into 384-well plates using an SH800S cell sorter (Sony Biotechnology).

### Bone marrow samples

Mononuclear cells were isolated from diagnostic BM samples using Ficoll-Paque (Cytiva) and viably cryopreserved in IMDM containing 20% FCS and 10% DMSO. For each sorting experiment, we included an aliquot of an adult BM sample as an internal control for flow cytometric analyses; this sample was obtained from the sternum of an adult patient undergoing cardiac surgery at the Academic Medical Center Amsterdam (AMC) after written informed consent was provided and was approved by the medical ethics review board of the AMC (MEC: 04/042#04 17,370). BM cells were thawed in IMDM containing 20% FCS, 100 µg/ml DNAse, and 5 mM MgCl_2_, resuspended in FACS buffer (PBS containing 0.5% FCS and 2 mM EDTA), and dissociated by filtering through a 70-µM cell strainer. Consecutive stains with the live/dead marker near-IR dye, surface markers, and DRAQ5 were performed at 4°C at a cell density of 10^7^ cells/ml (see **Table S6** for details). The cells were then sorted into 384-well plates using a FACSAria III cell sorter (BD Biosciences), cells were sorted into 384-well plates. We enriched for tumor cells by gating on CD34^neg^CD45^neg^CD90^pos^/GD2^pos^ cells (see **Fig. S1C**) after demonstrating that all but two tumor cells in a CD45^neg^-enriched fraction expressed CD90 and/or GD2 (see **Fig. S1D**). Mesenchymal surface markers (**Fig. S1B**) were included in order to discriminate between tumor cells and mesenchymal stromal cells (MSCs). We previously showed (*59*) that the CD34^neg^CD45^neg^CD90^pos^CD105^neg^ population in patient-derived BM samples contains tumor cells, identified using NB-specific qPCR(*14*).

After sorting, the plates were spun down, snap-frozen on dry ice, and stored at -80°C until further processing.

### Sequencing and processing of raw data

Library preparation was performed by scDiscoveries (Utrecht, Netherlands) using the SORT-seq protocol, a partially robotic version of the CEL-seq2 protocol (*24*). Paired-end 2×75-bp read length was used for sequencing on an Illumina NovaSeq6000 sequencer at the single-cell facility at the Princess Máxima Center (Utrecht, Netherlands) (for bone marrow samples) or on an Illumina NextSeq500 sequencer at the Hartwig Medical Foundation (Utrecht, Netherlands) (for the primary tumor samples). Raw data were processed using the Sharq pipeline (*65*). Read mapping was performed using STAR, version 2.6.1, on the hg38 Patch 10 human genome. Function featureCounts (RRID:SCR_012919) of the subread package (version 1.5.2) was used to assign reads based on GENCODE version 26 (RRID:SCR_014966). Generated count tables, per plate, were imported into R using Sharq helper functions in the SCutils package (*66*), per plate identifiers were added, and the count tables were then aggregated into one large count table for further analyses. A part of this scRNA-seq data has also been used in a joint study by Villalard *et al.* (manuscript under review).

### Quality control steps and normalization

Seurat version 4.3.0 (for quality control) and version 5.0.1 (for other analyses) were used for subsequent analysis. After removing spike-in genes, we filtered out cells with <500 or >45,000 transcripts, with >30% mitochondrial reads (see **Fig. S1E**), and/or high expression of hemoglobin. Ambient RNA was removed using the decontX function in the celda 1.8.1 package (*67*). The remaining 14,570 cells from 16 samples obtained from 9 patients were then normalized and scaled using either the NormalizeData and ScaleData functions or SCTransform in Seurat.

### Feature selection, dimensionality reduction, and cluster identification

Principal component (PC) analysis was performed on the 3,000 most variable genes, after removing confounders such as cell cycle, stress genes, cellular activity (i.e., ribosomal genes), sex (e.g., X chromosome-specific genes XIST and TSIX, and Y chromosome genes), and sample processing (see below). The first 50 PCs were then used for UMAP embedding. Clustering was performed using the FindNeighbors function (50 dimensions) and the FindClusters function.

### Distinguishing between malignant and non-malignant cells

To identify the cell types, we used a 3-step process. First, cell type labels were assigned using SingleR (*68*), comparing the gene expression patterns in clusters of interest with the pattern reported in a reference dataset (the Human Cell Atlas) (see **Fig. S1G**). Next, cells with neuronal characteristics were identified preliminarily as tumor cells, while neuronal cells with high expression levels of *SOX10* and *S100B* were identified as Schwann cells; similarly, we assessed the expression of cell type-specific genes for other cell types to support our cell type identification (see **Fig. 1C,E**). Finally, the malignancy of clusters assigned as tumor cells was confirmed by CNV analysis using inferCNV, with non-malignant cells (i.e., immune and endothelial cells) in our dataset serving as a reference (see **Fig. S1H**; immune cells from PT samples were further analyzed in Wienke *et al* (*69*)). The tumor cluster was further refined by repeating the above-mentioned steps. Tumor clusters consisting of 2,404 primary tumor cells and 4,446 DTCs were selected for further analysis.

### Correcting for differences in sample processing

In analyses in which we compared tumor cells in paired PT and BM samples, we used a correction step to account for differences in sample preparation (see “Sample processing” above). We performed differential gene expression analysis between PT and BM cell populations for: *i*) stromal cells, *ii*) T cells, *iii*) cytotoxic T cells, *iv*) NK cells, and *v*) endothelial cells (data not shown). The overlap of these DEG lists, filtered by *p*-value (*p*<0.05) and directionality (by log2FC), was saved as “processing-related genes” (**Table S7**). These genes are expected to capture differences between cells in the PT and BM samples that are due to technical artifacts (e.g., enzymatic digestion and/or thawing of cryopreserved cells) and not biological characteristics of the respective cell types; these genes were therefore removed from the list of the most variable genes that underlie dimension reduction and clustering.

### Differential gene expression and gene set enrichment analysis

Differential gene expression (DGE) was performed using the FindMarkers function in Seurat. The RNA assay of the Seurat object was used, and ribosomal genes were excluded. For PT vs BM analyses, DGE was performed on all 7 PT samples vs. the corresponding 7 BM samples, while correcting for patient differences. For GD2-high vs GD2-neg/low analyses, DGE was performed using cells of patient 1+2 in each group, the two patients who had the largest variation in GD2 expression. To compare between PT and BM samples (**Table S1**), and between GD2-high and GD2-neg/low DTCs (**Table S5**), GSEA was performed using MsigDB (*70*) with a false discovery rate (FDR) *q*-value of <0.05. Gene set sources included Gene Ontology (GO), HALLMARK, REACTOME, KEGG, and TFT (transcription factor targets). For comparing between PT and BM samples, DEGs with log2FC>|0.5| and an adjusted *p*-value of <0.05 were used as input for GSEA (n=117 and 120 PT and BM cells, respectively). To compare between GD2-high and GD2-low DTCs, DEGs with log2FC>|1| and an adjusted *p*-value of <0.05 were used as input for GSEA (n=500 and 57 GD2-high and GD2-low cells, respectively). GSEA to compare between Chr9-normal and Chr9-loss tumor cells in Patient 2 (**Table S2**) was performed using the fgsea R-package (*71*), using the complete DEG list as input, and gene sets were subsequently filtered (Benjamini-Hochberg-adjusted *p*-value of <0.05).

### CNV heterogeneity

CNV heterogeneity between paired sets of PT and BM samples in each patient was determined by assessing the inferCNV output plots and determining chromosomes (or regions of chromosomes) that showed a distinctly different pattern for the PT compared to BM (or vice versa). The inferCNV package was run as described previously (*28*), using the following parameters: cutoff=0.3, denoise=TRUE, and sd_amplifier=2. In addition, we re-labeled each potential tumor cluster using the source of the cell, introducing a combined patient identifier and biopsy type variable (PT or BM). This enabled us to assess the CNVs per patient and biopsy type. Chromosome 9 aberration status per cell for Patient 2 was defined using CONICS R package as described previously (*72*).

### Whole-genome sequencing

CNVs were detected from WGS data for the primary tumor as part of the diagnostics workflow at the Princess Máxima Center. Details regarding sample preparation, quality control, and CNV identification were published previously by Van Belzen *et al*. (*73*). In short, WGS libraries were sequenced using the NovaSeq 6000 sequencing platform (Illumina), and WGS data were analyzed in accordance with the Princess Máxima Center standardized pipelines using GATK 4.0 Best Practices (*74*).

### Expression of gene sets

The expression of gene lists was assessed using the Seurat-internal function AddModuleScore; with the exception of the comparison between PT and BM cells, in which a manual score was created per cell by calculating the mean expression of genes from the respective gene set, using the scaled expression and thereby correcting for sample processing (see above).

### Cell cycle and drug sensitivity scoring

To identify cells as either high cycling or low cycling, we used G2M-phase and S-phase scores available within Seurat (see **Fig. S2B**). To predict drug sensitivity, lists of genes for which the expression was associated with sensitivity to drugs were downloaded from the DepMap portal (*75*) and filtered for significance (defined as an adjusted *p*-value of <0.05) and the slope of the correlation. A gene was assigned to the sensitivity score if that gene’s expression was correlated negatively with the area under the curve derived from the dataset’s drug screen. A list of genes in the sensitivity score for each drug is provided in **Table S3**.

### Dormancy signature

Gene signatures containing genes that are differentially expressed in dormant cells compared to proliferating tumor cells published in two previous studies (*31*, *32*) were filtered (logFC>|1| and *p*<0.05), and the overlap between the genes with positive log2FC (i.e., upregulated in dormant cells) or negative log2FC (i.e., downregulated in dormant cells and upregulated in proliferating cells) was used as the final dormancy and proliferation signatures, respectively (**Table S4**). Expression of this signature in DTCs was then assessed using the AddModuleScore function in Seurat.

### Data integration

Data from 9 BM samples was integrated for analysis of BM-DTCs (see **Figs. 4 and 5**), after SCTransform normalization as described previously (*76*).

### Index analysis

BM cells were FACS index sorted; that is the process of recording flow cytometric data for single cells that are sorted into a plate and simultaneously assigning a well ID (“index”), which is used in subsequent analyses for integrating flow cytometry data with sequencing data. To this end, Flow Cytometry Standard (FCS) files were imported in R using the FlowCore package (*77*), and then combined into a single cell object using the CATALYST R package (*78*). Data were transformed using arcsinh transformation with cofactor 150 and added to the Seurat object based on the well ID (“index”).

Thresholds for cell surface expression intensity (negative, low, medium, or high expression) were established using non-malignant cell populations (stromal, B, T, and NK cells) as reference (see **Fig. S5E**).

### Determination of subcellular localization for gene lists

Ensemble IDs of genes were used as input in the IDmapping tool of the UniProt database (https://www.uniprot.org/id-mapping/). The “Subcellular location [CC]” column was included in the output and served as the variable for selecting genes whose protein products were annotated as localized to the plasma membrane based on experimental evidence (ECO:0000269), in addition to genes that are known to encode cell adhesion molecules or receptors (**Table S5**).

### NB cell lines and NB tumoroids

The high-risk NB cell lines (n=23) included 13 MYCN-amplified and 10 non-*MYCN*-amplified lines. *MYCN*-amplified: SKNBE, SJNB6, SJNB8, NGP, NMB, SJNB10, IMR32, TR14, CHP134, KPNYN, KCNR, LAN5 and N206. Non-*MYCN*-amplified: SKNSH, SKNAS, SKNFI, SY5Y, SKNMM, LAN6, SHEP2, GIMEN, SJNB12 and SJNB1. Cells were grown in culture flasks at 37°C in humidified air containing 5% CO2, and were passaged every 4-7 days. Culture medium consisted of DMEM high glucose (ThermoFisher Scientfic, #11965092) with 10% FBS (Sigma-Adrich, #F0804), 5 mM L-Glutamine (ThermoFisher Scientific, #25030024), 1x MEM non-essential amino acids solutions (ThermoFisher Scientific, #11140035) and 500 U/ml penicillin-streptomycin (ThermoFisher Scientific, #15140122).

Tumoroids 691-ADRN and 691-MES (*79*) were grown in culture flasks at 37°C in humidified air containing 5% CO_2_, and were passaged every 4-7 days by mechanical dissociation (691-ADRN) or with StemPro Accutase Cell Dissociation Reagent (Gibco, #A1110501) (691-MES). The culture medium consisted of Dulbecco’s modified Eagle’s medium (DMEM) with low glucose, Glutamax supplement and pyruvate (Gibco, #10567014), 25% Ham’s F-12 Nutrient Mix (Gibco, #11765054), B-27 supplement minus vitamin-A (Life Technologies, #12587010), N-2 supplement (Life Technologies, #7502048), 1% penicillin-streptomycin (Life Technologies, #15140122), 20 ng/ml animal-free recombinant human EGF (PeproTech, #AF-100-15), 40 ng/ml recombinant human FGF-basic (PeproTech, #100-18B), 200 ng/ml recombinant human IGF-I (PeproTech, #100-11), 10 ng/ml recombinant human PDGF-AA (PeproTech, #100-13A), and 10 ng/ml recombinant human PDGF-BB (PeproTech, #100-14B). All cell lines and tumoroids were authenticated by STR profiling and were routinely checked for mycoplasma contamination.

### Transcriptomics and lipidomics analyses of NB cell lines

For the transcriptomic and lipidomic analyses, the 23 NB cell lines were cultured in 6-well plates in triplicate. The media was refreshed 24 hours before harvesting. Transcriptomics was performed by USEQ (Utrecht, the Netherlands). The library preparation was performed using poly-AAA capture. Paired-end 2x 50 bp read length was used for sequencing on Illumina NextSeq2000 sequencer. The raw data was processed using RNA-SEQ pipeline version 2.0. Readcounts were then generated using the Subread FeatureCounts module (v2.0.3) with the Homo_sapiens.GRCh38.106.ncbi.gtf file as annotation, after which normalization was done using the R-package edgeR (v3.40). For transcriptomic analysis of NB tumoroids, a dataset containing Affymetrix microarray gene expression data from ADRN and MES pairs of tumoroids (GSE90803) was accessed using the “R2: Genomics Analysis and Visualization Platform” (*80*) and assessed for *NGFR* mRNA levels.

Lipidomic analysis was conducted at the Core Facility Metabolomics, Amsterdam UMC (Amsterdam, the Netherlands), following established protocols as previously described (*81*). Briefly, after addition of internal standards, lipid extraction was performed using one-phase chloroform/methanol extraction and this was analyzed using a Thermo Scientific Ultimate 3000 UPLC (with both normal phase and reversed phase separations) coupled to a Q-Exactive Plus Orbitrap mass spectrometer. Mass spectrometry data were collected in both positive and negative ionization modes and analyzed with an in-house lipidomics pipeline written in the R programming language (http://ww.r-project.org) and MATLAB. Lipid identification was based on accurate mass, retention times, and fragmentation spectra, with semi-quantitative results normalized to internal standards. The threshold for categorizing GD2 lipid levels into low, medium, and high groups was based on the 33rd percentile. Lipid values below this threshold (<1.9) were classified as low. Group classifications were further validated at the protein level by flow cytometry analysis of GD2 on a selection of 5 cell lines, ensuring consistency between lipid-based and protein-based groupings.

### Flow cytometry of NB tumoroids

Tumoroids were dissociated into single cells and stained with a LIVE/DEAD™ Fixable Near-IR Dead Cell Stain Kit (Thermo Fisher, #L34975) for 15 min and subsequently with antibodies against NGFR and GD2 for 20 minutes at 4°C (**Table S6**). Surface expression was measured using on a ID7000 spectral cell analyzer (Sony Biotechnology) and analyzed using the manufacturer’s software.

### Statistics

Differences in gene (signature) expression were determined using the Wilcoxon rank sum/ Mann-Whitney U test (in the case of 2 groups), or the Kruskal-Wallis rank sum test (in the case of 3 or more groups) followed by Dunn’s test for multiple comparisons. Spearman correlation was used to analyze the correlation between gene expression and surface marker expression and was assessed using the Spearman coefficient (ρ) and *p*-value.

## List of abbreviations

ADRN: adrenergic
BM: bone marrow
CNV: copy number variation
DEG: differentially expressed gene
DGE: differential gene expression
DTC: disseminated tumor cell
EMT: epithelial-to-mesenchymal transition
GSEA: gene set enrichment analysis
ITH: intra-tumor heterogeneity
MES: mesenchymal
MNA: MYCN amplification
PT: primary tumor
scRNA-seq: single-cell RNA sequencing
UMAP: uniform manifold approximation and projection

## Acknowledgments

We thank Philip Lijnzaad, Thanasis Margaritis, and Aleksandra Balwierz at the single-cell genomics facility of the Princess Máxima Center for support. Illustrations in figures were created with BioRender.com.

## Funding

KiKa, grant number 374 (IT, CH, GAMT)

Sanquin Research grant for Product and Process development for Cellular products, number 19-27 (IT, CV)

Princess Máxima Center grant number P0104 (IT, CH, GAMT)

Innovative Medicines Initiative 2 Join Undertaking, grant agreement No 116064, https://www.itccp4.eu (AB, KMK, SvH, JJM)

European Research Council (ERC) under the European Union’s Horizon 2020

Research and Innovation Program, grant agreement number 716079 Predict (AB, KMK, SvH, JJM)

EraCoSysMed grant INFER-NB (AB, KMK, SvH, JJM)

Stichting Villa Joep—The Neuroblastoma Immuno-consortium (AB, KMK, SvH, JJM)

Stichting Sijmen (AB, KMK, SvH, JJM)

These sponsors had no involvement in the study design or the collection, analysis, or interpretation of the data.

## Author contributions

Conceptualization: CH, AB, KMK, SvH, CV, JJM, GAMT, IT

Methodology: CH, AB, KMK, ST, SvH, CV, JJM, GAMT, IT

Investigation: CH, KMK, ST, MvdM, IT

Formal analysis: CH, AB, KMK, ST, MvdM, CPB, ZvGF, SvH, IT

Data curation: CH, AB, KMK, CPB, ZvGF

Visualization: CH, AB, CBP, IT

Software: AB, SvH

Resources: ABPvK, CV, JJM, GAMT, IT

Project administration: JJM, GAMT, IT

Supervision: ABPvK, SvH, CV, JJM, GAMT, IT

Funding acquisition: JJM, GAMT, IT

Writing—original draft: : CH, AB, KMK, MvdM, JJM, GAMT, IT

Writing—review & editing: CH, AB, KMK, CPB, ZvGF, SvH, CV, JJM, GAMT, IT

## Data and materials availability

All data are available in the main text or the supplementary materials. The datasets generated for and analyzed in this study are available in the Gene Expression Omnibus repository and are accessible through GSE245175.

## Supplementary Materials

Fig. S1-S5 and Tables S1-S7 are provided as separate files.

