## Supplementary Figures 1-5 for "Metastatic tumor cells in bone marrow differ from paired neuroblastoma tumor and contain subsets with therapy-resistant characteristics"

**This PDF file includes:**

Fig. S1 to S5

**Other Supplementary Materials for this manuscript include the following:**

Table S1 to S7

### Supplementary Figure 1

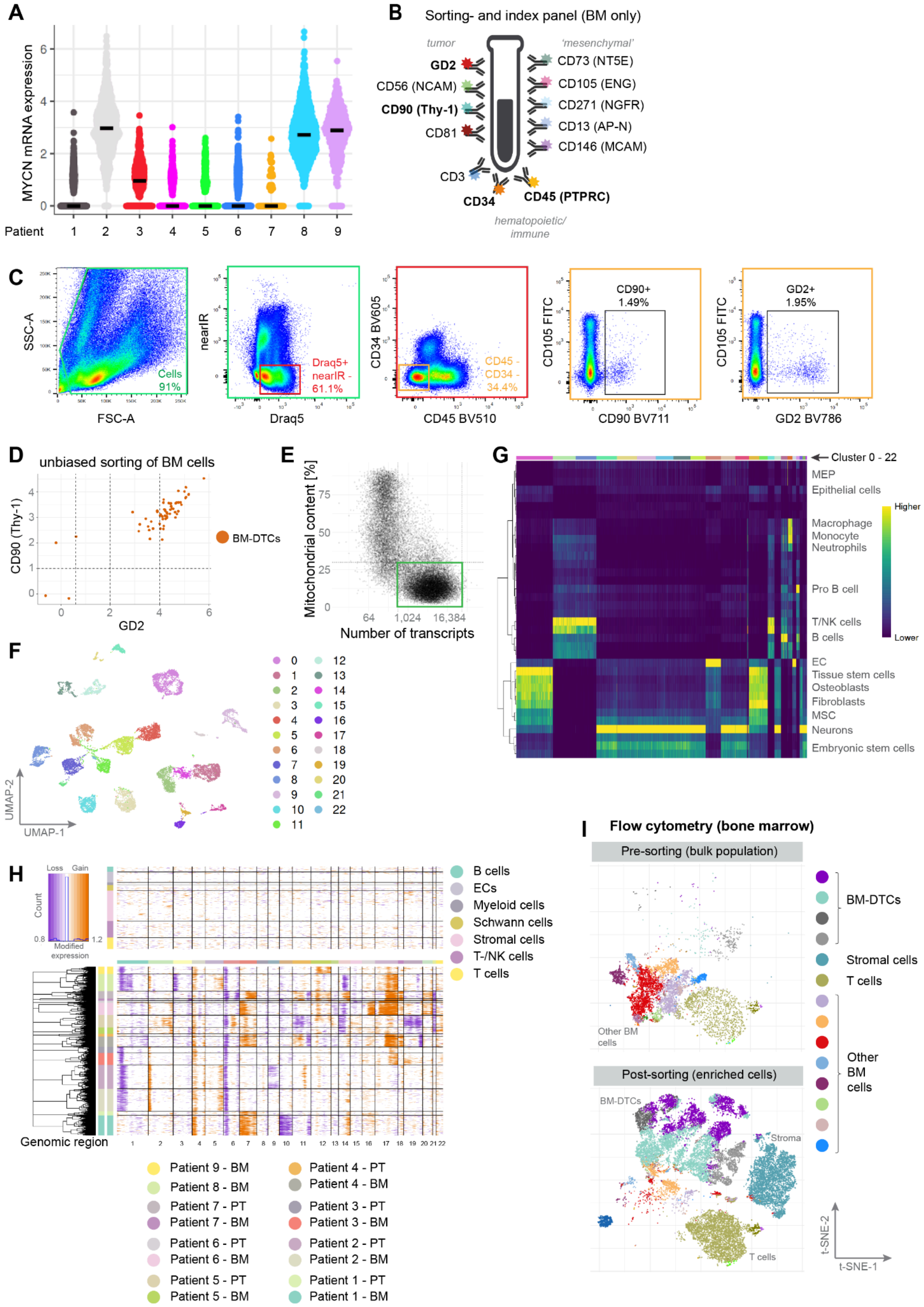

**Fig. S1. Index sorting strategy and cell type identification. A)** Violin plot of *MYCN* mRNA levels measured in the tumor cells for all 9 patients. The thick horizontal lines represent the median value. **B)** The panel of 12 surface markers used for sorting and index analysis of BM samples. The four markers indicated in bold were used to enrich for tumor cells. Mesenchymal markers were included in order to discriminate between tumor cells and mesenchymal stromal cells. **C)** Representative FACS gating strategy used for enriching for DTCs (shown is an example from patient 5). The CD34-CD45- population (orange) was gated on CD90+ /GD2+ cells. The percentage of the parent gate is indicated for each gate. **D)** CD90 expression plotted against GD2 expression in DTCs from patient 3 (identified using sequencing data) that were sorted using an unbiased approach (i.e., enriching only for CD45- cells). Each symbol represents a single tumor cell. Note that all but two tumor cells expressed CD90 and/or GD2. **E)** The thresholds for quality control of sequencing data. Cells with <500 transcripts and/or >25% mitochondrial content were excluded from further analyses. **F)** UMAP of all cells in the dataset (including non-malignant cells), annotated as 23 clusters. **G)** Heatmap showing the resemblance of each of the 23 clusters to known cell types in the Human Cell Atlas using SingleR algorithm. **H)** CNV analysis for all 9 patients using the inferCNV algorithm. The expression intensity across each chromosome is compared between clusters of interest (lower panel) and non-malignant reference clusters (upper panel). Purple and orange indicate a chromosomal loss and gain, respectively. **I)** Dimension reduction (t-SNE) with flow cytometry data for all BM samples. The FCS (Flow Cytometry Standard) files of all recorded cells ("bulk", top panel) and the FCS files of all 384-well plates (i.e., the sorted and sequenced cells, bottom panel) were used as input. Mesenchymal stromal cells and T cells were intentionally enriched for, but were not analyzed in this study.

**Supplementary Figure 2**

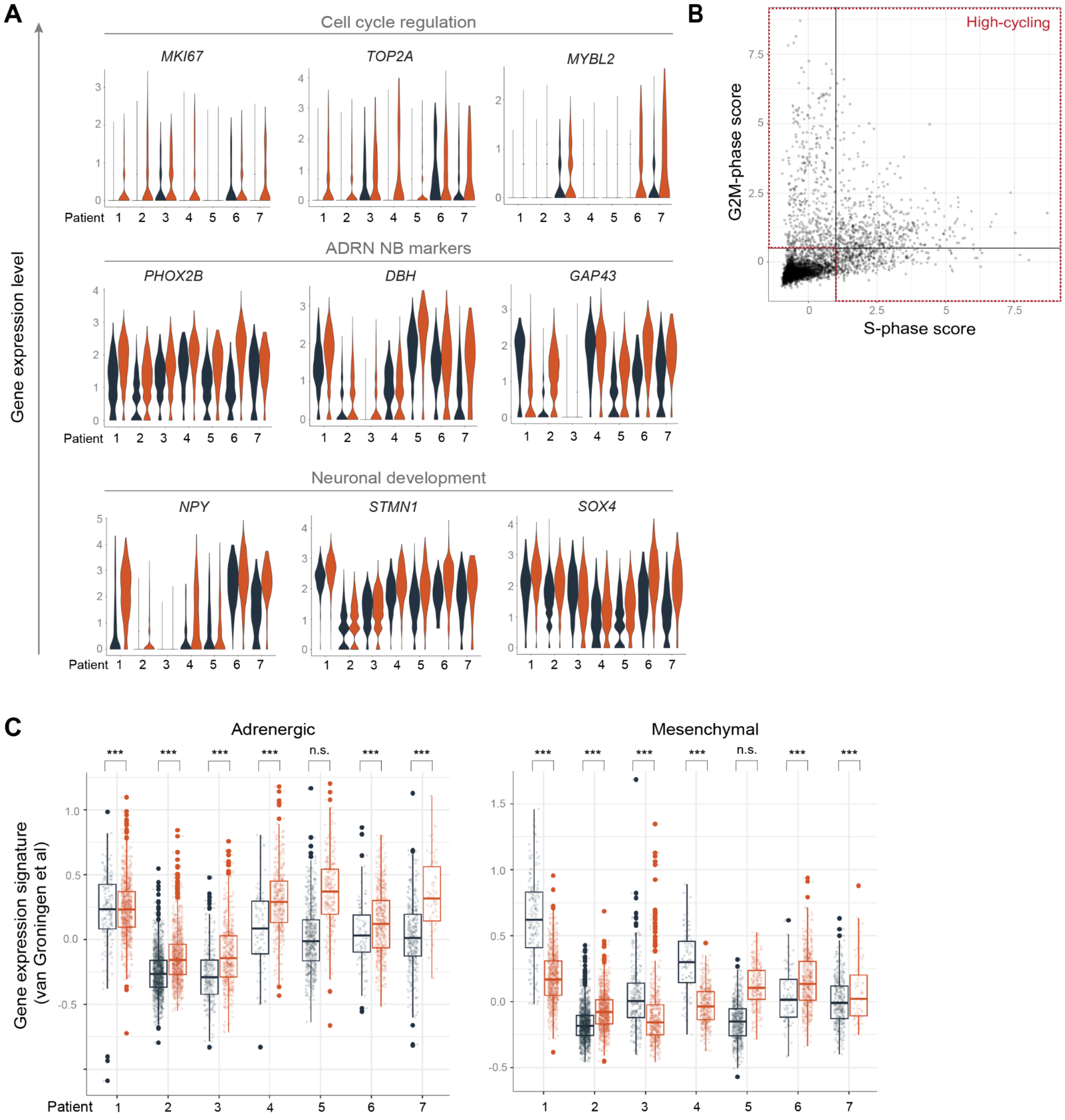


**Fig. S2. Difference in transcriptomics between tumor cells from the primary tumor and paired DTCs in the bone marrow. A)** Violin plots showing the expression levels of select differentially expressed genes in PT cells and BM-DTCs, per patient (see Table S1 for the full list). **B)** Thresholds for defining "non-cycling" (bottom-left) and "high-cycling" (top-right) categories (used in Fig. 2E) based on expression of S-phase and G2M-phase gene expression scores. **C)** Box plots summarizing the expression of alternative adrenergic and mesenchymal gene signatures (derived from Van Groningen *et al*, 2019) (12) in BM and PT sample, per patient. Gene expression was assessed on scaled data in order to correct for processing differences between PT and BM samples (see Methods). ****p*<0.001 and n.s., not significant (Wilcoxon rank sum test).

#### Supplementary Figure 3A

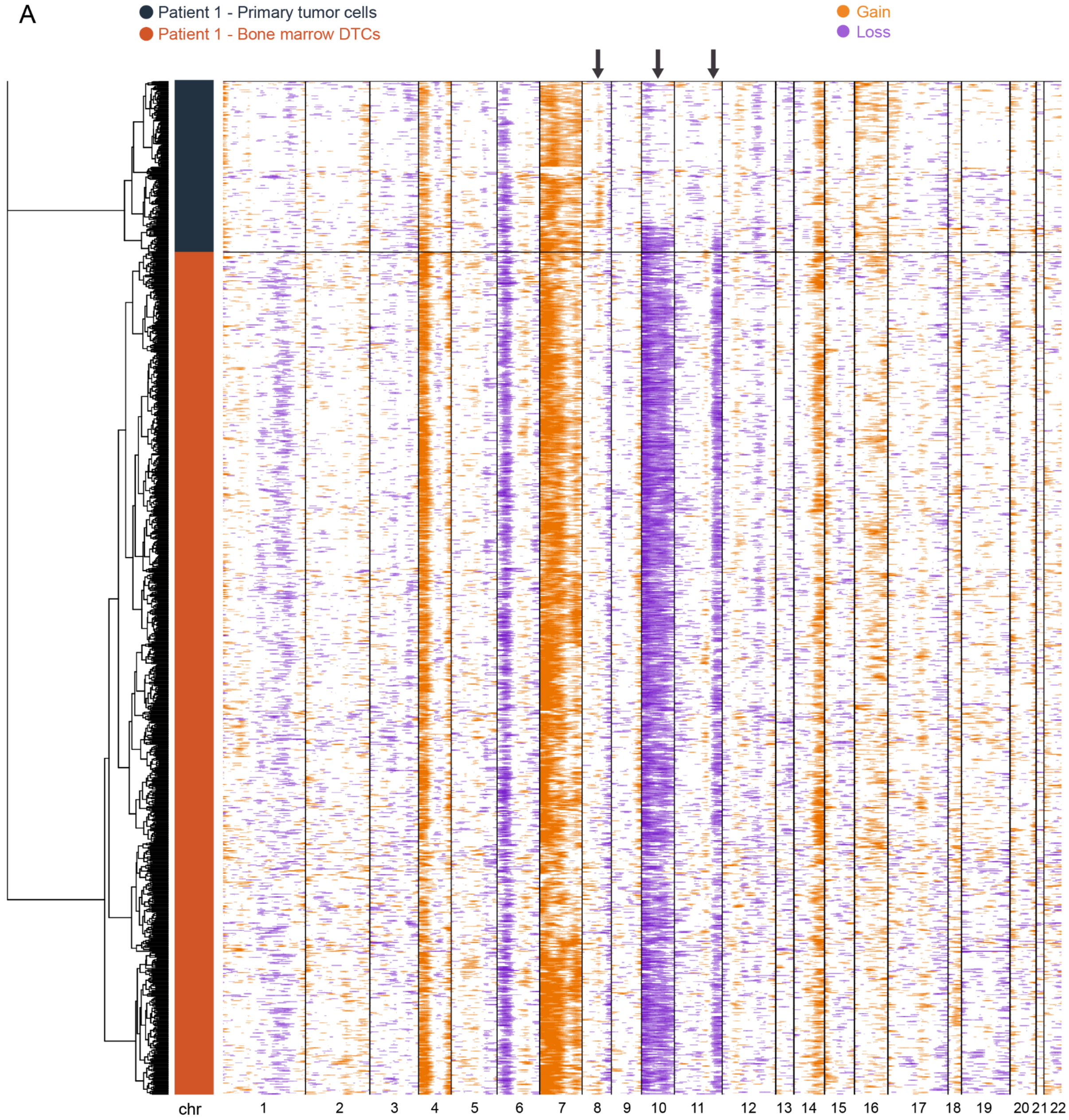

**Supplementary Figure 3B-C**

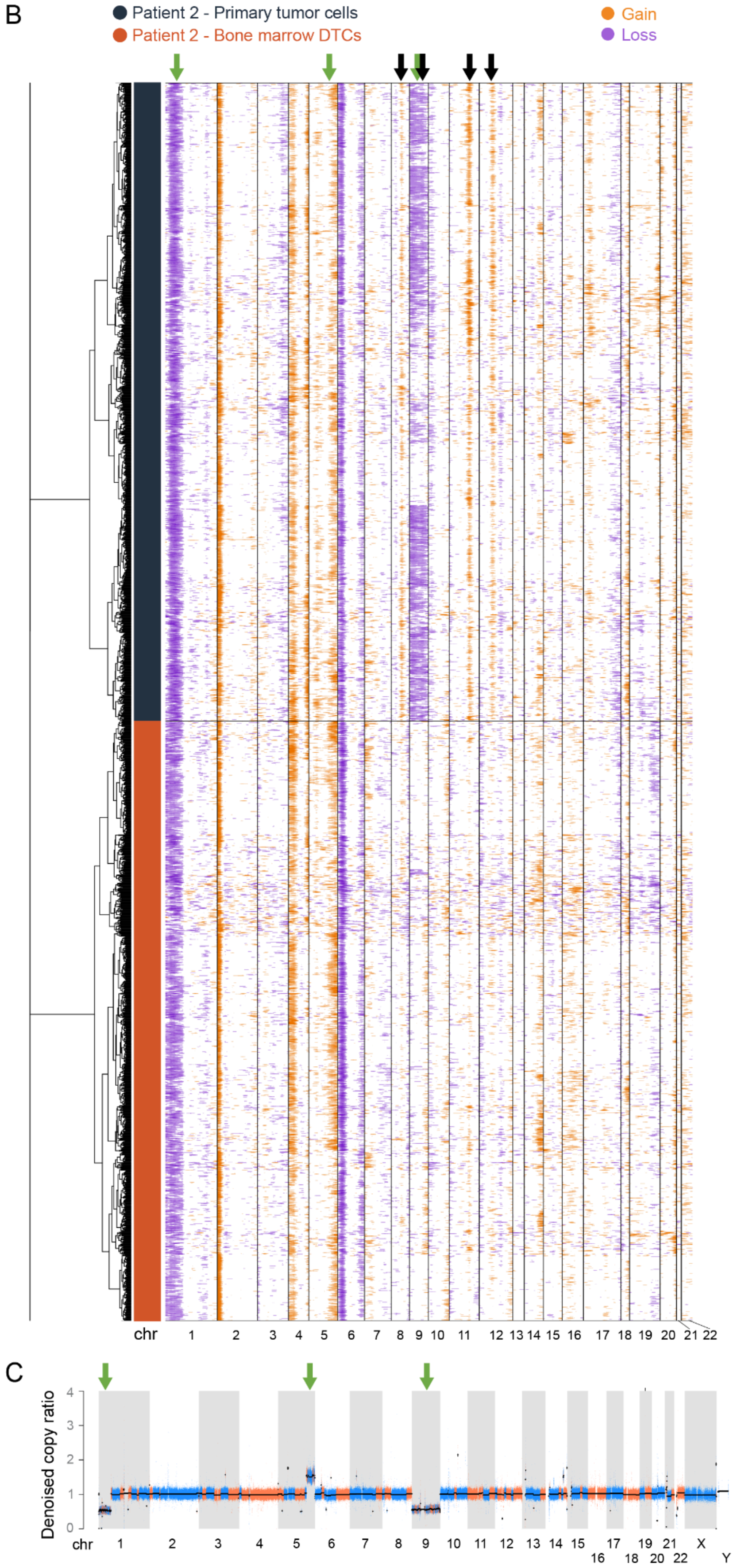

**Supplementary Figure 3D-E**

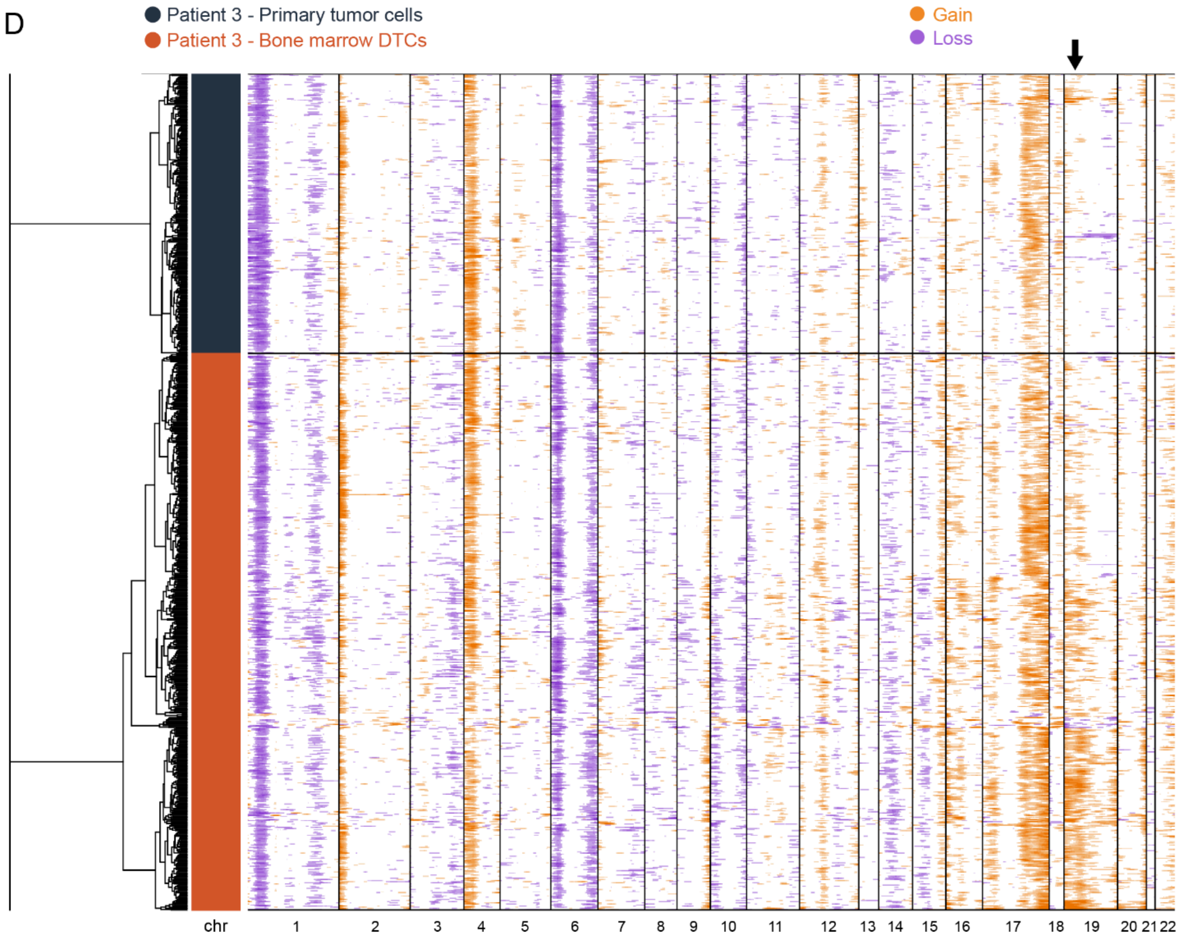


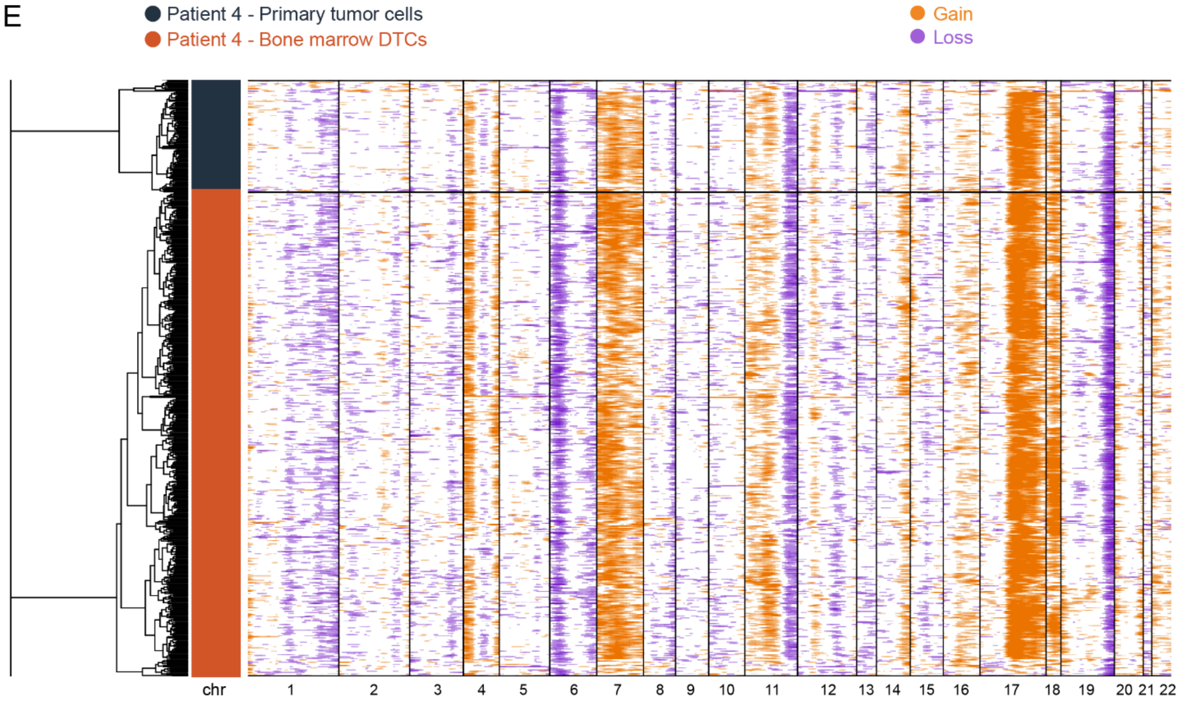

**Supplementary Figure 3F-G**

F

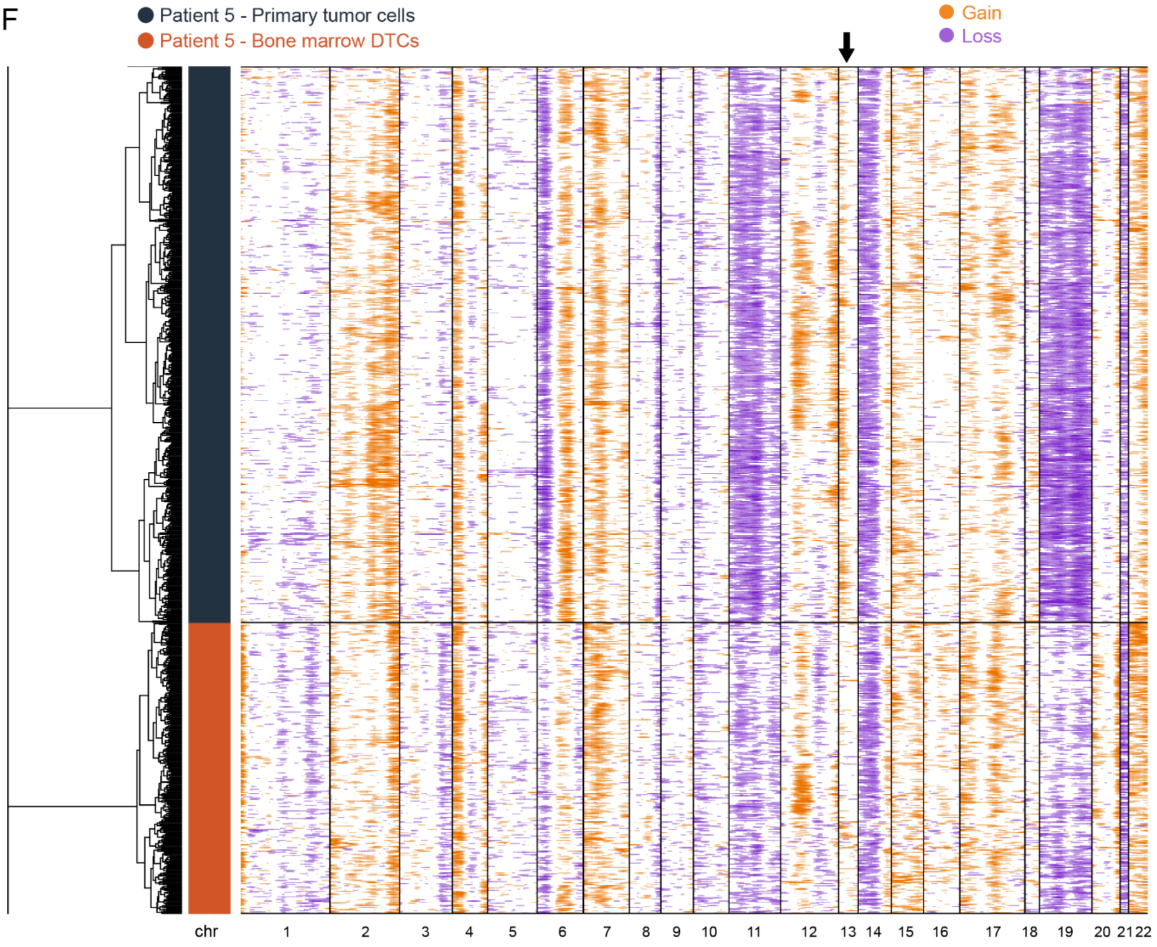


G

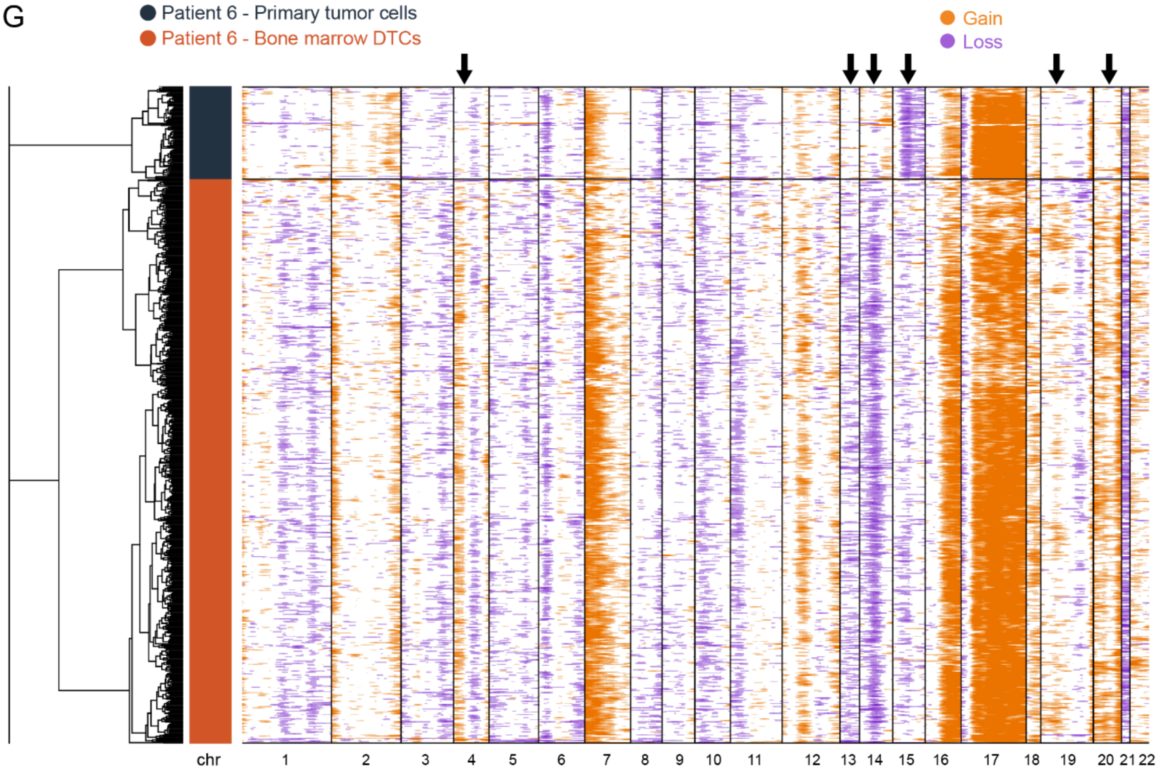

**Supplementary Figure 3H**

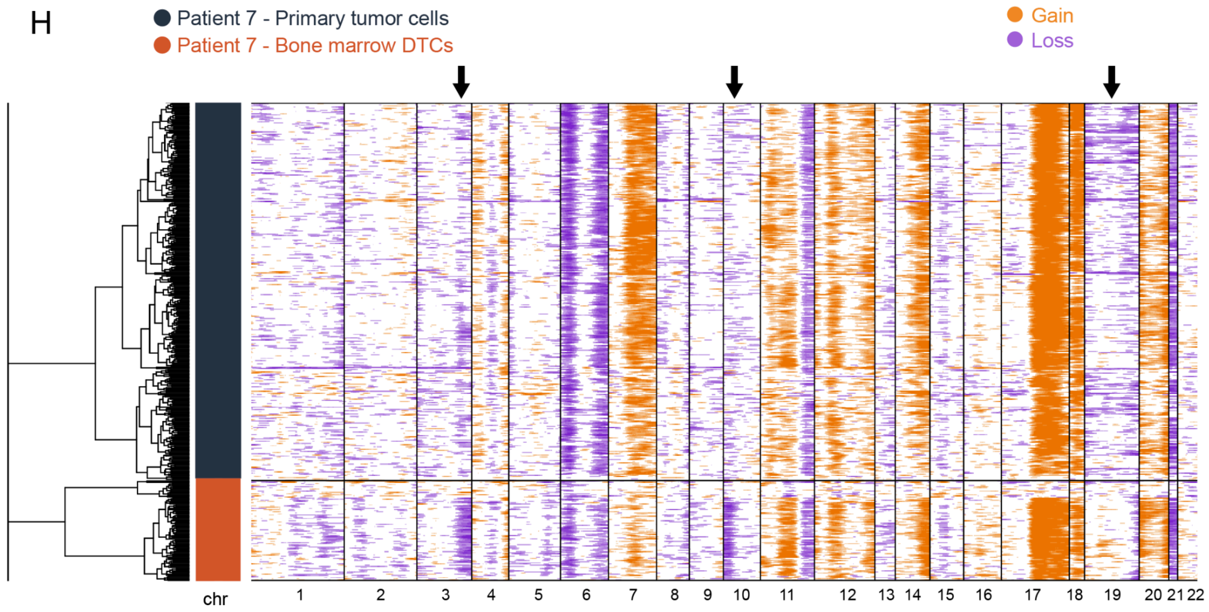


**Fig. S3. Assessment of copy number variations in PT and BM, for all 22 somatic chromosomes per patient**. Shown are detailed CNV maps (produced using CONICS) to discern heterogeneity in CNVs between PT cells and DTCs. Black arrows indicate the chromosomes with patterns of aberrations that differ between PT and DTCs. Aberrations indicated from clinical diagnostic analyses (i.e., whole-exome sequencing) are provided below for each patient. Note that CNVs routinely tested for are located in chromosomes 1p, 11q, 17q. **A)** Patient 1 (clinical diagnostics: none). **B)** Patient 2 (clinical diagnostics: *MYCN* amplification, 1p loss, and 11q gain); see Fig. 3B for a more condensed view of this patient's data. Green arrows indicate the aberrations that were also detected using whole-genome sequencing of the PT of this patient. **C)** Whole-genome sequencing data of the PT in Patient 2, showing a loss in chromosome 1p, amplification in chromosome 5q, and a deletion across the entire chromosome 9. These aberrations (green arrows) and additional ones were also detected using the RNA-based CNV algorithm in (B). **D)** Patient 3 (clinical diagnostics: *MYCN* amplification, 1p partial loss, and 17q partial gain). **E)** Patient 4 (clinical diagnostics: 11q partial loss, 11q partial gain, and 17q partial gain). **F)** Patient 5 (clinical diagnostics: 17q gain). **G)** Patient 6 (clinical diagnostics: multiple copies of chromosome 2, 1p gain, and 17q gain). **H)** Patient 7 (clinical diagnostics: 1p partial loss, 11q partial loss. 11q partial gain, and 17q partial gain).

#### Supplementary Figure 4

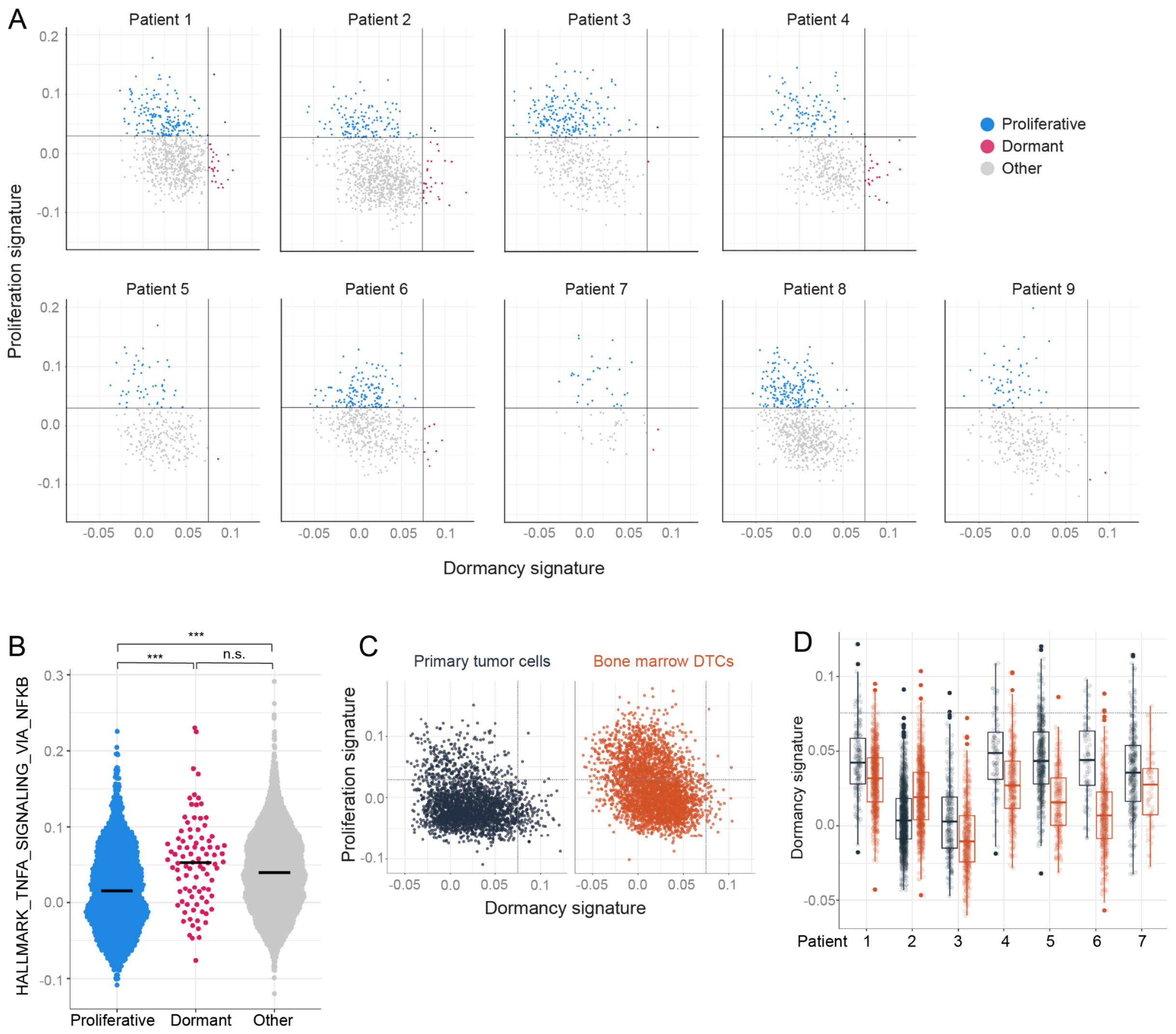


**Fig. S4. Further characterization of dormancy in DTCs in all patients and in the primary tumor cells. A)** Expression of gene signatures characterizing DTCs as either dormant, or proliferating or neither of the two in each patient. **B)** Violin plot summarizing the expression levels of the HALLMARK gene set "TNF-α signaling via NF-κB" in dormant and proliferating cells (see Fig. 4D for definitions of dormancy and proliferation). ***$p$<0.001 and n.s., not significant (Kruskal-Wallis rank sum test followed by Dunn's test for multiple comparisons). **C)** Expression of gene signatures characterizing dormant and proliferating cells in all PT cells and DTCs obtained from Patients 1-7. **D)** Box plot summarizing the expression of the dormancy signature in PT cells and DTCs for Patients 1-7. Boxes indicate the inter-quartile range, and the thick horizontal lines indicate the median value. The thresholds are copied from Fig. 4D.

**Supplementary Figure 5**

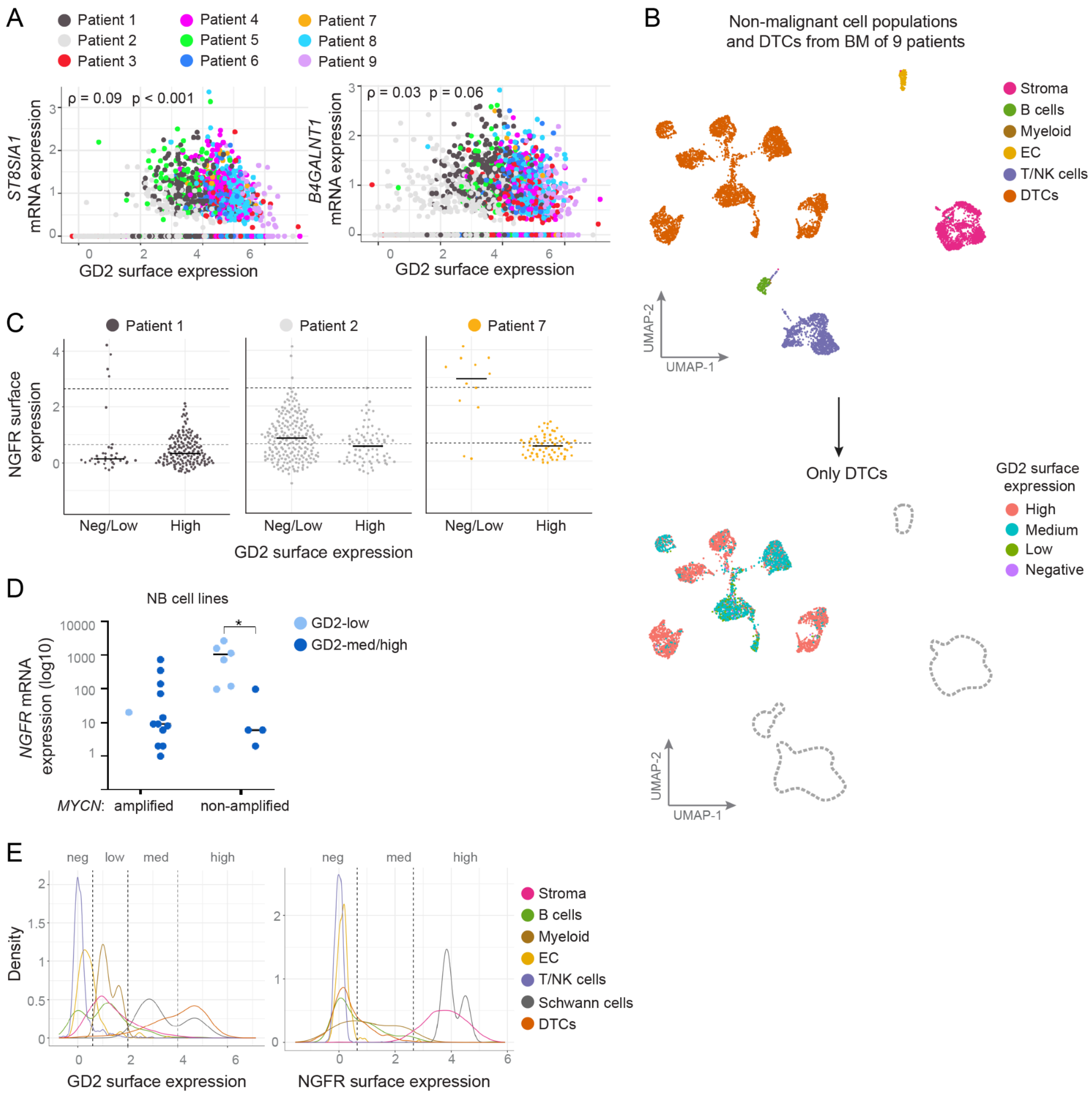


**Fig. S5. Cell surface protein expression in DTCs. A)** The mRNA levels of the genes encoding the two main enzymes directly involved in GD2 synthesis plotted against GD2 surface expression. Spearman coefficients: ρ=0.09 and ρ=0.03 for *ST8SIA1* and *B4GALNT1*, respectively; *p*<0.001 and *p*=0.06 for *ST8SIA1* and *B4GALNT1*, respectively. **B)** UMAP of non-malignant and malignant BM cells that were sorted from the samples of 9 patients. Tumor cells from all four GD2 categories (see Fig. 5A-B) exclusively align with tumor clusters, and do not cluster in the UMAP space of non-malignant cell types. **C)** NGFR surface expression measured in GD2-low/neg and GD2-high DTCs in BM samples from 3 patients, based on a threshold of ≥10 GD2-low DTCs. Surface expression was assessed per cell by flow cytometry during cell sorting. **D)** *NGFR* mRNA levels measured in 23 NB cell lines (Fig. 5E), divided into *MYCN*-amplified (n=13) and non-amplified (n=10) groups. GD2 levels were determined by lipidomics. The thick horizontal lines indicate the median value; and the *y*-axis is a log scale. Experiments were performed in triplicate. *$p<0.05$ (Kruskal-Wallis test). Note that the majority of *MYCN*-amplified cell lines are GD2-med/high. **E)** Density plot of normalized GD2 (left) and NGFR (right) surface expression measured in DTCs and non-malignant cell populations for Patients 1-9. The thresholds indicate GD2 expression (negative, low, medium, and high) and were set based on the population distribution in malignant and non-malignant cell types. EC, endothelial cell, NK, natural killer.

**Other Supplementary Materials (separate files)**

**Table S1**. Differentially expressed genes and gene set enrichment analysis for comparison between bone marrow disseminated tumor cells and primary tumor cells

**Table S2.** Differentially expressed genes and gene set enrichment analysis for comparison between primary tumor cells of patient 2 with and without a chromosome 9 loss

**Table S3.** Drug sensitivity scores per cluster of DTCs and per dormancy-categories, and list of genes included in each sensitivity score

**Table S4.** List of genes in dormancy signature

**Table S5.** Differentially expressed genes and gene set enrichment analysis for comparison between disseminated tumor cells with high and low GD2 surface expression

**Table S6.** List of antibodies used for index sorting of bone marrow samples

**Table S7.** List of genes used for correction of sample processing
